# *In vivo* glucose imaging in multiple model organisms with an engineered single-wavelength sensor

**DOI:** 10.1101/571422

**Authors:** Jacob P. Keller, Jonathan S. Marvin, Haluk Lacin, William C. Lemon, Jamien Shea, Soomin Kim, Richard T. Lee, Minoru Koyama, Philipp J. Keller, Loren L. Looger

## Abstract

Glucose is arguably the most important molecule in metabolism, and its mismanagement underlies diseases of vast societal import, most notably diabetes. Although glucose-related metabolism has been the subject of intense study for over a century, tools to track glucose in living organisms with high spatio-temporal resolution are lacking. We describe the engineering of a family of genetically encoded glucose sensors with high signal-to-noise ratio, fast kinetics and affinities varying over four orders of magnitude (1 µM to 10 mM). The sensors allow rigorous mechanistic characterization of glucose transporters expressed in cultured cells with high spatial and temporal resolution. Imaging of neuron/glia co-cultures revealed ∼3-fold higher glucose changes in astrocytes versus neurons. In larval *Drosophila* central nervous system explants, imaging of intracellular neuronal glucose suggested a novel rostro-caudal transport pathway in the ventral nerve cord neuropil, with paradoxically slower uptake into the peripheral cell bodies and brain lobes. In living zebrafish, expected glucose-related physiological sequelae of insulin and epinephrine treatments were directly visualized in real time. Additionally, spontaneous muscle twitches induced glucose uptake in muscle, and sensory- and pharmacological perturbations gave rise to large but enigmatic changes in the brain. These sensors will enable myriad experiments, most notably rapid, high-resolution imaging of glucose influx, efflux, and metabolism in behaving animals.

## Introduction

D-Glucose is the central molecule of energy metabolism in the cycle of life. It is a preferred source of energy for most bacteria^1^ and fungi, the photosynthetic product of plants, and the primary circulating source of energy in animals. Dysregulation of glucose is the hallmark of diabetes, and is also central to obesity and metabolic syndrome^2^. Together, these conditions contribute to a burgeoning global health crisis, and exacerbate most other health problems, including cancer^3^ and heart disease^4^, among others. Tools to monitor glucose are thus of paramount importance to both clinical diagnoses and basic science. In the brain, there are many mysteries surrounding glucose in particular and bioenergetics in general. Glucose is the primary energy source of the brain, but in what cells does glucose metabolism occur? Do glycolysis and oxidative phosphorylation^5^ occur in the same cells? Do neurons in fact utilize glucose directly or is it first converted to lactate by the proposed astrocyte-neuron lactate shuttle^6^? Do synaptic terminals rely on local glycolysis to support their energy-intensive processes^7^? What are the roles of specific glucose transporters^8^, *e.g.* Glut1, Glut3 and Glut4, in neuronal and glial glucose uptake?

To address these and other critical questions about glucose import, export, synthesis, consumption, *etc.*, tools that can specifically detect glucose are required. The most useful tools will satisfy the following criteria: they should 1) be easily targetable to specific cell types, sub-cellular compartments and organelles, and even to specific proteins of interest such as glucose transporters; 2) be genetically encoded (*i.e.* protein-based) to enable generation of viruses and transgenic animals; 3) be soluble proteins (as opposed to integral membrane proteins) to provide maximal flexibility in targeting, such as to the cytoplasm or inside specific organelles; 4) be compatible with long-term imaging, and stable over time, to facilitate longitudinal studies; 5) not require any cofactors, other added molecules, or endogenous molecules of non-uniform distribution; 6) provide for cellular and sub-cellular resolution detection; 7) respond roughly as fast as the underlying phenomena; 8) not perturb physiological phenomena being measured, and 9) ideally be amenable to straightforward detection without sophisticated equipment.

Techniques for glucose monitoring in fluids (blood, urine), mainly to assist diabetic patients in maintaining euglycemia, have existed for decades and have gone through multiple cycles of optimization^9^. Current methods for *ex vivo* glucose monitoring, however, are incompatible with the demands for *in vivo* detection listed above. The dominant technology for *ex vivo* glucose monitoring is indirect detection of glucose levels through electrochemical detection of H_2_O_2_ release from glucose breakdown by the enzyme glucose oxidase^10^. This technique is sensitive and rapid, but incompatible with our needs: electrochemical detection reports from only single sites^11^ and lacks cellular resolution and targetability. The primary method of *in vivo* glucose tracking (used usually as a proxy for metabolic activity) is systemic injection of the non-metabolizable, radiolabeled glucose analogue 2-[F^18^]fluoro-2-deoxy-D-glucose (FDG), whose accumulation is imaged by position emission tomography (PET)^12^. FDG, however, is not targetable, PET’s spatial resolution is fundamentally limited to several millimeters^13^, and each scan takes from tens of seconds to minutes. Since FDG is not metabolized, it has dramatically different pharmacokinetics from glucose, and as such is only an approximation of initial glucose uptake. Thus, it is clear that these methods do not fit the criteria mentioned above, and better methods for *in vivo* glucose monitoring are required.

Fluorescence microscopy has revolutionized the study of physiology, allowing high spatiotemporal resolution of signaling events in intact animals and plants. The use of genetically encoded indicators further allows imaging to be targeted to specific cell types of interest and even to sub-cellular compartments; stable expression through transgenics, viral transduction and other methods allows longitudinal experiments up to years. The first fluorescent glucose sensors were developed from the plant lectin concanavalin A^14^. These sensors, however, required *in vitro* derivatization with small-molecule dyes and injection, and fluorescence changes were either poor (*in vitro*) or unmeasurable (plasma)^15^. Specificity was also lacking, since concanavalin A also binds to other sugars and glycans^16^.

The cloning of bacterial periplasmic glucose-binding proteins has led to a wide array of sensing modalities. The introduction of single cysteine residues allows coupling of thiol-reactive small-molecule fluorophores that respond to glucose binding with alterations in fluorescence intensity, lifetime, emission wavelength, or FRET ratios^17 18,19^, but such sensors would require injection into living specimens, with unknown targeting and poor *in vivo* stability. The first fully genetically encoded fluorescent glucose sensor resulted from the fusion of cyan and yellow fluorescent proteins (CFP, YFP) to the termini of *Escherichia coli* glucose/galactose-binding protein (GGBP)^20^. These initial sensors showed modest response amplitudes (<10% change in fluorescence resonance energy transfer (FRET) between CFP-YFP), but allowed imaging of glucose influx into cultured mammalian cells. Iterative structure-based optimization of these sensors increased FRET change to ∼60%^21^, which facilitated experiments such as identification of the SWEET family of glucose transporters mediating pollen viability and bacterial virulence/control, among other functions^22^, and glucose signaling in rice roots in response to diverse stressors^23^.

Despite the advances in this family of indicators (fluorescent indicator proteins for glucose, FLIPglu), the modest fluorescence responses have hampered application to preparations such as living animals. Our lab has recently developed a technique for engineering single-wavelength indicators from circularly permuted green fluorescent protein (cpGFP) and periplasmic binding proteins for a variety of small molecules: glutamate^24,25^, GABA^26^, maltose^27^, phosphonates^28^, among others. Another group has also used this technique to develop sensors for histidine^29^ and recently glucose (sensor “FGBP” made using *E. coli* GGBP)^30^. FGBP was specific for D-glucose and D-galactose over other sugars tested, and produced a ∼200% change in fluorescence excitation ratio in *E. coli* cells upon addition of glucose. However, the FGBP sensor was not tested in any other preparation.

Here we present a series of genetically encoded green glucose sensors with high signal-to-noise ratio, rapid kinetics and simple imaging modality. We also demonstrate use of these sensors in a number of applications: characterization of glucose transporter function and inhibitor properties, separate visualization of neuronal and glial metabolic rates in co-culture, discovery of a novel putative glucose transport pathway in fruit fly brains, and tracking of glucose transport in the muscles and brains of living larval zebrafish. These sensors will facilitate longitudinal glucose imaging studies in living animals and plants, with high spatial and temporal resolution.

## Results

### Sensor engineering and protein characterization

Similar to the designs of several other genetically encoded small molecule sensors based on bacterial periplasmic binding proteins (PBPs), circularly permuted green fluorescent protein (cpGFP) was inserted into an appropriate glucose-binding PBP (GBP) to create a sensor. A survey of the Protein Data Bank revealed several structures of GBP’s from multiple organisms, including mesophiles *E. coli, Pseudomonas putida, Yersinia pestis, Salmonella typhimurium* and *Chloroflexus aurantiacus*, and hyperthermophiles *Thermus thermophilus* and *Thermotoga maritima*. Because proteins from thermophiles often exhibit enhanced stability, the GBP from the thermophile *T. thermophilus* was selected for a scaffold. It also had high structural similarity to *E. coli* maltose-binding protein, from which we previously made a sensor^27^, thus facilitating engineering.

A gene encoding the *T. thermophilus* GBP lacking the periplasmic secretion leader peptide (amino acids 1-26) was synthesized and cloned into bacterial expression vector pRSET (Invitrogen). Based on homology to the maltose sensor, GlyGly-cpGFP-GlyGly was inserted between residues 326 and 327 of TtGBP (**Fig. 1a-b**). Residues His66 and His348, which form aromatic stacking interactions with bound glucose, were mutated to alanine to reduce glucose affinity for screening purposes. Both GlyGly linkers were randomized by targeted mutagenesis, and variants screened for high changes in fluorescence upon addition of saturating glucose. A variant, Linker1-ProAla/Linker2-AsnPro (L1-PA/L2-NP) was identified with ΔF/F = 2.3 and K_d_ for glucose of 2.3 mM. The additional mutation Leu276Val was made to further decrease affinity to 7.7 mM, while preserving ΔF/F (**Fig. 1e**). Stopped-flow fluorescence showed relatively rapid rates of signal increase upon binding glucose (k_on_ = 31 M^-1^s^-1^) and signal decrease upon glucose unbinding (k_off_ = 0.38 s^-1^) (**Fig. 1f**). We refer to this sensor as intensity-based glucose-sensing fluorescent reporter (iGlucoSnFR; not to be confused with our other sensor iGluSnFR, which responds to glutamate). We tested the binding of iGlucoSnFR to a range of sugars (**Supp. Fig. 1**); only D-galactose and 2-deoxy-glucose responded at all, with much weaker affinity and lower ΔF/F. iGlucoSnFR responded to glucose across a range of pH values from 6.0-9.5 (**Supp. Fig. 2**), with K_d_ and ΔF/F fairly similar, but both trending higher with increased pH, across the range 7.0-9.5.

**Figure 1:**
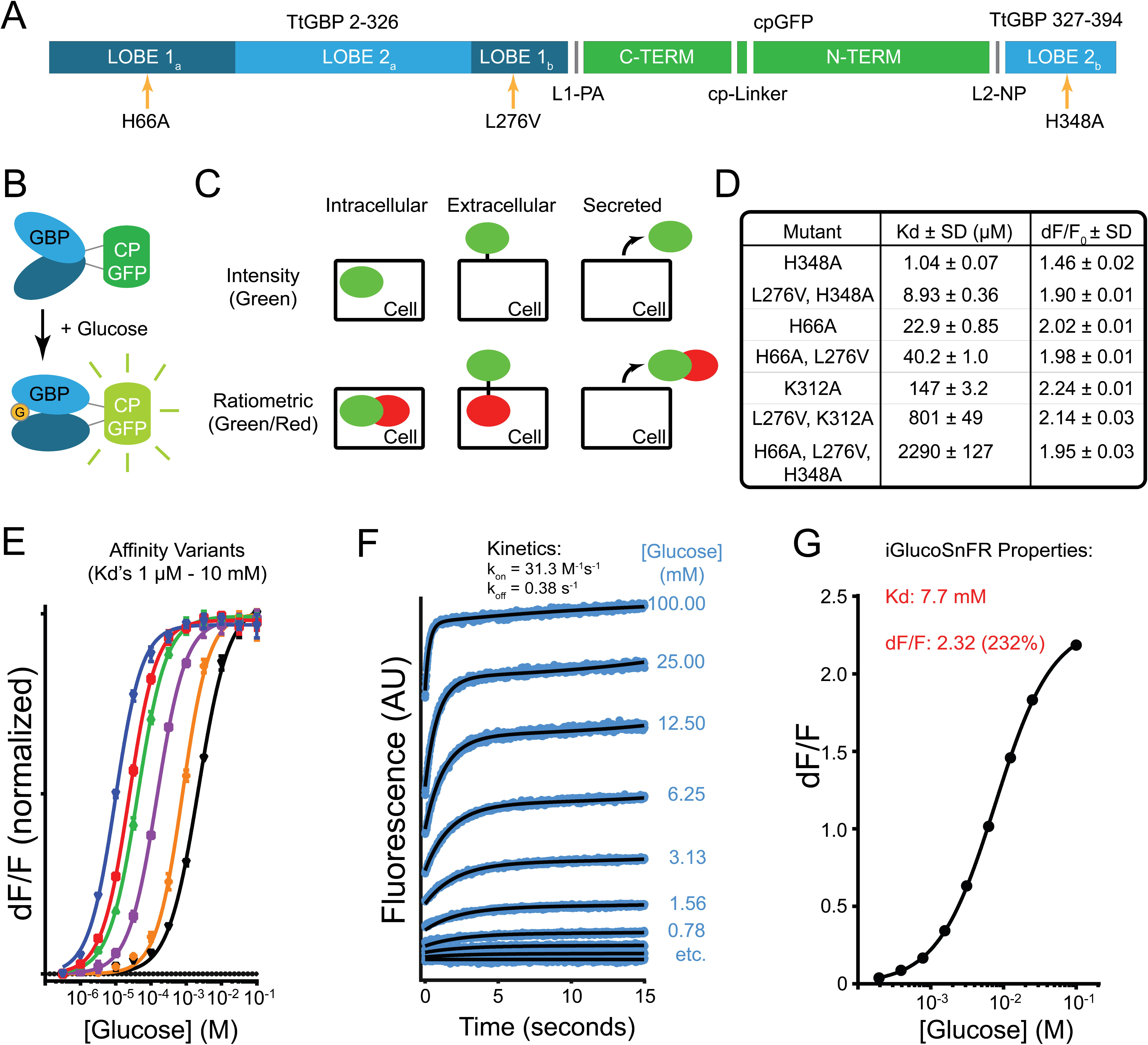
Sensor Design and Properties. A: Schematic of iGlucoSnFR sensor sequence. Circularly-permuted green fluorescent protein (cpGFP, green) was inserted into glucose-binding protein (GBP, light and dark blue) from *T. thermophilus.* Affinity-lowering mutations are indicated (orange arrows). B: Schematic of iGlucoSnFR sensor construction. Circularly-permuted green fluorescent protein (cpGFP, green) was inserted into glucose-binding protein (GBP, light and dark blue) from *T. thermophilus* such that binding of glucose (G, orange disk) to GBP induced fluorescence increases in cpGFP. C: Sensor architectures. Sensors were made that reside intracellularly, are attached to the extracellular membrane, or are secreted from the expressing cell. Similar sensors were also made with the addition of a glucose-insensitive red fluorescent protein mRuby2 to allow for ratiometric measurements and/or cell typing. Sensors are shown schematically as green ellipses, mRuby2 as red ellipses. D,E: Affinity variants. Permutations of binding site residue mutations (H66A, L276V, H348A) yielded iGlucoSnFR variants with affinities spanning the range of 1 µM to 10 mM. Each curve represents a sensor with different affinity, with fluorescence changes normalized. F: iGlucoSnFR kinetics. Stopped-flow fluorescence measurements on the final construct were performed at several concentrations, and kinetic rate constants were fit to data. G: Final iGlucoSnFR binding curve, constructed from the data in (F). Sensor parameters as shown. Error bars (standard deviations) are plotted, but are smaller than the data points.

For use in eukaryotic cells, we cloned iGlucoSnFR, without the N-terminal affinity purification sequence from the bacterial expression vector, into a *CMV* promoter-containing vector derived from pDisplay (Invitrogen) for cytoplasmic expression (**Fig. 1c**). Although not discussed here, we also created vectors for membrane surface display of iGlucoSnFR (pDisplay) or secreted in soluble form (by removing the transmembrane domain from pDisplay). Altogether, we created six versions: the green sensor alone or with the red fluorescent protein mRuby2^31^ fused to the C-terminus (see below), each in the three targeting formats. Titration of purified iGlucoSnFR-mRuby2 showed sensor properties slightly shifted in K_d_ relative to the untagged sensor and with an apparently reduced ΔF/F (**Supp. Fig. 3**). The bacterially-expressed fusion, however, showed evidence of degradation on an SDS-PAGE gel, so it is difficult to ascribe reliability to these *in vitro* measurements. Moreover, since the tagged- and untagged sensors responded identically to identical stimuli in cell culture experiments described below, it is likely that the apparently reduced ΔF/F is an artifact due to the above-mentioned difficulties of purifying homogeneous fusion protein.

Finally, permutations of the three different binding site mutations (H66A, L276V, H348A) were tested to produce an affinity series of six sensors spanning four orders of magnitude (**Fig. 1d-e**).

### Imaging glucose transporter activity

We have previously demonstrated the utility of cytoplasmic sensors to report the activity of membrane transporters in cultured cells with excellent spatial (sub-cellular) and temporal (milliseconds) resolution^32^. HEK293T cells were co-transfected with plasmids encoding cytoplasmic iGlucoSnFR and the human glucose uniporter Glut1/ SLC2A1^33^. Buffers alternating between 0 and 20 mM glucose (substituted with equimolar sorbitol to maintain constant osmolarity) (**Fig. 2a**) were perfused continuously onto cells, with a periodicity of 60 seconds. Corresponding fluorescence changes were imaged on an inverted Zeiss LSM 800 confocal microscope at 2 Hz; ∼10 oscillation periods were measured to establish baseline oscillation magnitudes and variability. The Glut1 inhibitor cytochalasin B (CCB, a competitive inhibitor) was then superadded to these buffers at varying concentrations. CCB reversibly decreased Glut1 transport activity in a dose-dependent manner (**Fig. 2b**). CCB binds to the intracellular side of Glut1^34^ and thus must enter the cell before it can act. Consistent with that mechanism, steady-state inhibition was achieved after about two oscillation periods at each concentration. Once steady-state inhibition was achieved, magnitudes were quantified relative to baseline, and an IC_50_ of ∼0.6 µM was fit to these data, consistent with literature values^34^.

**Figure 2:**
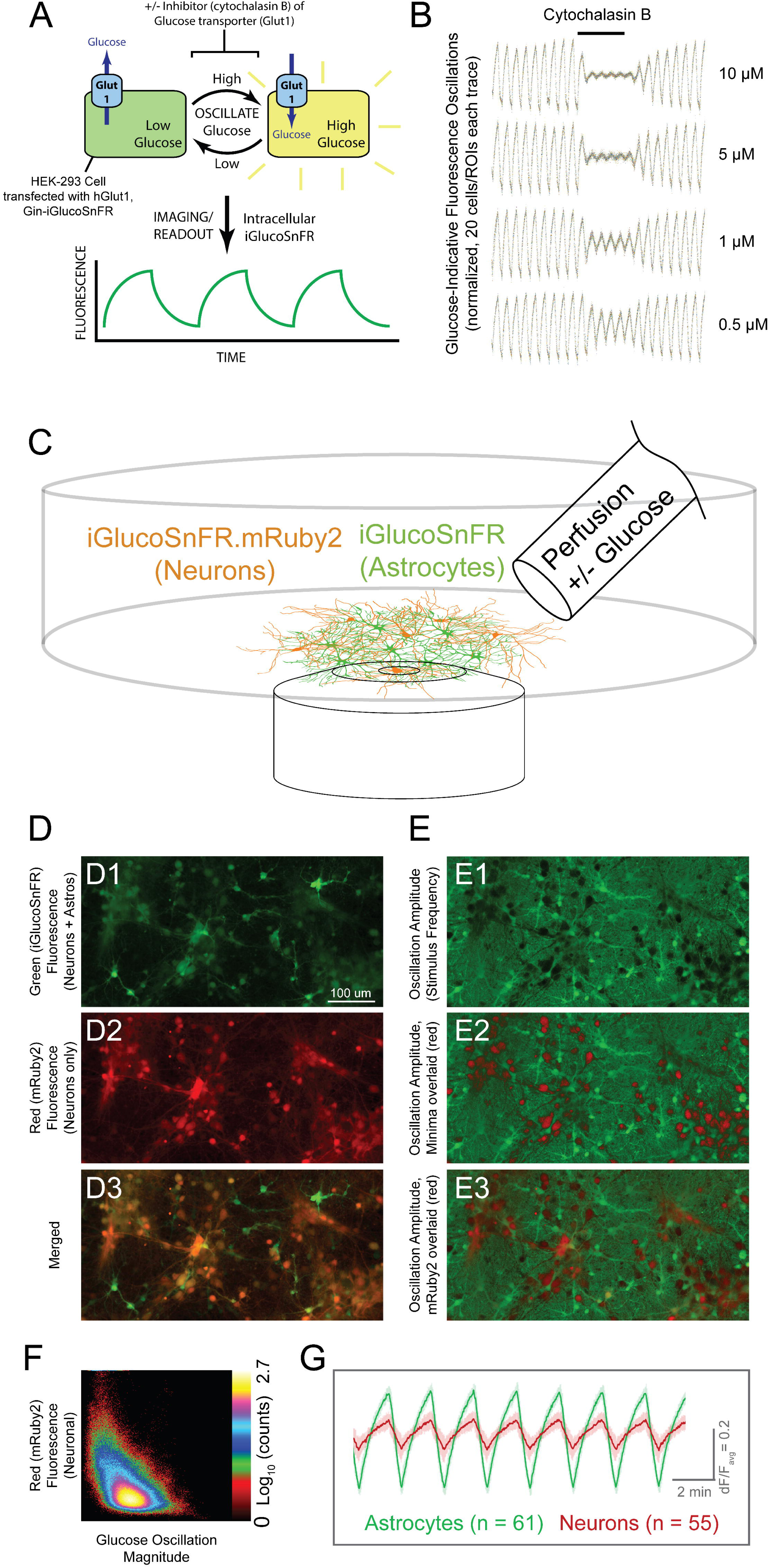
Sensor Applications in Cell Culture. A: Schematic of transporter assay. Cultured cells (HEK293) were co-transfected with plasmids for expression of iGlucoSnFR and the human glucose uniporter hGlut1. Buffers were oscillated +/-glucose, eliciting oscillating fluorescence responses, which were subsequently diminished by overlaying constant concentrations of the glucose transporter inhibitor cytochalasin B at various concentrations. B: Oscillating fluorescence responses and inhibition. Twenty regions of interest (ROIs) corresponding to cells were identified by fast Fourier transform (FFT) magnitude in initial segments without inhibitor, and were filtered over the whole experiment by subtraction of a moving-window-average of length corresponding to the stimulus period. Each trace was then normalized by RMSD during the pre-treatment phase and plotted as individual data points. In the middle segment (indicated at top), cytochalasin B was added at the concentrations indicated at the right of each trace. After removing cytochalasin B, oscillation magnitudes returned to initial values. Imaging parameters were: two channels: Ex 488 nm, Em 505–550; Ex 561, Em 565–650; objective EC Plan-Neofluar 10×/0.3 NA M27, imaging rate ∼1 Hz; 512 x 512 pixels; stimulus period 60 s. C: Schematic of neuronal culture assay. Acute astrocyte-neuron co-cultures from rat hippocampus were infected with adeno-associated viruses driving iGlucoSnFR-mRuby2 expression in neurons (*hSynapsin1* promoter) and green-only iGlucoSnFR in astrocytes (*gfaABC*_*1*_*D* promoter). ACSF + 20 mM glucose or sorbitol was continuously perfused, and fluorescence changes measured. D: Fluorescence imaging of co-cultures. (D1-3) fluorescence images for each channel are shown as indicated. (E1-3) Images of pixelwise temporal FFT magnitudes at stimulus frequency, indicative of glucose concentration changes. E1 shows amplitudes in green. E2 shows the same image but with FFT minima overlaid in red, for comparison with the mRuby2 signals overlaid in E3 (also see alone in D2). Two channels: Ex 488 nm, Em 505–550; Ex 561, Em 565–650; objective Plan-Apochromat 20x/0.8 M27, imaging rate ∼1 Hz; 1024 x 512 pixels; stimulus period 120 s. E: Correlation plot between FFT magnitude (x-axis) and mRuby2 signal (y-axis). Note clear (albeit weak) anti-correlation between the two (CC ∼-0.5): high intensity in the red channel corresponds to weak magnitudes and vice versa. G: Averaged fluorescence responses (ΔF/F_avg_) to oscillating stimulus from astrocytes (identified by high-thresholded green:red fluorescence ratio) and neurons (identified by high red fluorescence signals). ΔF/F_avg_ was calculated by subtraction of a moving window average with size corresponding the period of the stimulus oscillations followed by division by the same moving average. Note larger magnitude of glucose response in astrocytes. Error bars represent standard deviations in fluorescence responses within each group. Nearly identical results with reversed tags are shown in **Supp. Fig. 4**.

### Characterization of glucose dynamics in neuronal/glial co-cultures

Next, we measured intracellular responses of co-cultured rat astrocytes and neurons to approximately physiological levels of glucose. Adeno-associated viruses (AAVs) were used to express iGlucoSnFR and iGlucoSnFR-mRuby2 under promoters specific for neurons (*hSynapsin1*) or astrocytes (*gfaABC*_*1*_*D*). Cultures were co-infected with AAV2/1-*hSynapsin1*-iGlucoSnFR-mRuby2 and AAV2/1-*gfaABC*_*1*_*D*-iGlucoSnFR, thus labeling both cell types with the glucose sensor and neurons with an additional red fluorescent protein (**Fig. 2c**). These cultures were subjected to continuously-perfused buffers alternating between 0 and 20 mM glucose with a total period of two minutes, with relative times in each buffer adjusted to preserve approximate linearity in fluorescence response waveforms (80 seconds with glucose, 40 seconds without). Constant confocal imaging in both green and red channels (**Fig. 2d, Supp. Movies 1a-b**) revealed ∼3-fold larger fluorescence oscillations in astrocytes compared to neurons (**Fig. 2e-g**). Oscillation magnitudes were nearly identical regardless of which cell type expressed the mRuby2-fused sensor (**Supp. Fig. 4**). Although the cell body-free background (**Fig. 2e1**) appears to have significant oscillation amplitude, close inspection reveals that this background is a dense network of cellular processes that are enhanced by fast Fourier transform (FFT) processing and the ΔF/F_avg_ calculation. The oscillation patterns of these processes correlate better with the green-only (astrocytic) fluorescence signal (**Fig. 2d1**) than with the green + red (neuronal) signal (**Fig. 2d2**). The astrocyte cell bodies have roughly similar amplitudes as the processes’ amplitudes, indicating that the processes are likely astrocytic. In this experiment, red fluorescence intensity correlated inversely with oscillation magnitude, providing a holistic picture of the relative indifference of neurons to glucose in comparison to that of astrocytes (compare **Fig. 2e1&2;** correlation is plotted in **Fig. 2f**).

Although metabolic consumption of glucose contributes modestly to these fluorescence changes^35^, the magnitude of observed fluorescence changes indicates significantly larger glucose transport in astrocytes compared to neurons under these *in vitro* conditions, consistent with some previous results^36^ but in conflict with others^37^. (See supplementary material appendix for a discussion of transport versus metabolism estimations.) It is possible in the latter case that the absence of transport blockers in wash solutions allowed radiolabeled glucose analogues to escape prior to scintillation counting, skewing results. Further use of iGlucoSnFR, especially in more physiologically significant *in vivo* preparations, may be able to resolve these discrepancies, but at a minimum, the efficacy and utility of iGlucoSnFR in cultured preparations is demonstrated by the current data.

### Characterization in larval fly central nervous system explant

To demonstrate the utility of the sensor in intact systems, we created a transgenic UAS-iGlucoSnFR-mRuby2 *Drosophila* line. After balancing, this fly line was crossed with the *57C10*-GAL4 driver line^38^ to establish pan-neuronal expression. Whole central nervous systems (CNS) (including brain lobes and the ventral nerve cord, VNC) were excised from third instar larvae and mounted in a thin layer of 1% agarose. Buffers with or without 20 mM glucose were oscillated with a periodicity of 10 minutes (**Fig. 3a**). Confocal volumetric time-lapse imaging (8 x 5 µm planes, 1024 x 512 pixels, ∼9s/volume) of these brains (**Supp. Movies 2a-d**) showed increases in glucose concentration throughout the whole explant, with much greater changes in the VNC than in the brain lobes (**Fig. 3b**). Glucose concentrations (**Fig. 3c**) rose most quickly in the rostral VNC neuropil (and some large nerves – although since they have been severed by the dissection, this is likely non-physiological). A wave of glucose sensor response moved caudally within the VNC neuropil, and eventually outward to the peripheral neuronal cell bodies. The rostro-caudal peak-to-peak delay in the neuropil was about 30 seconds. Influx into the brain lobes was delayed relative to the rostral VNC, and appeared similarly to proceed from the inside outwards. These results were reproducible across individual larvae (**Fig. 3d;** n = 5) and were independent of perfusion direction. Observation of spatiotemporal patterns of glucose trafficking such as this are currently only possible through the use of iGlucoSnFR or other fluorescent sensors like it.

**Figure 3:**
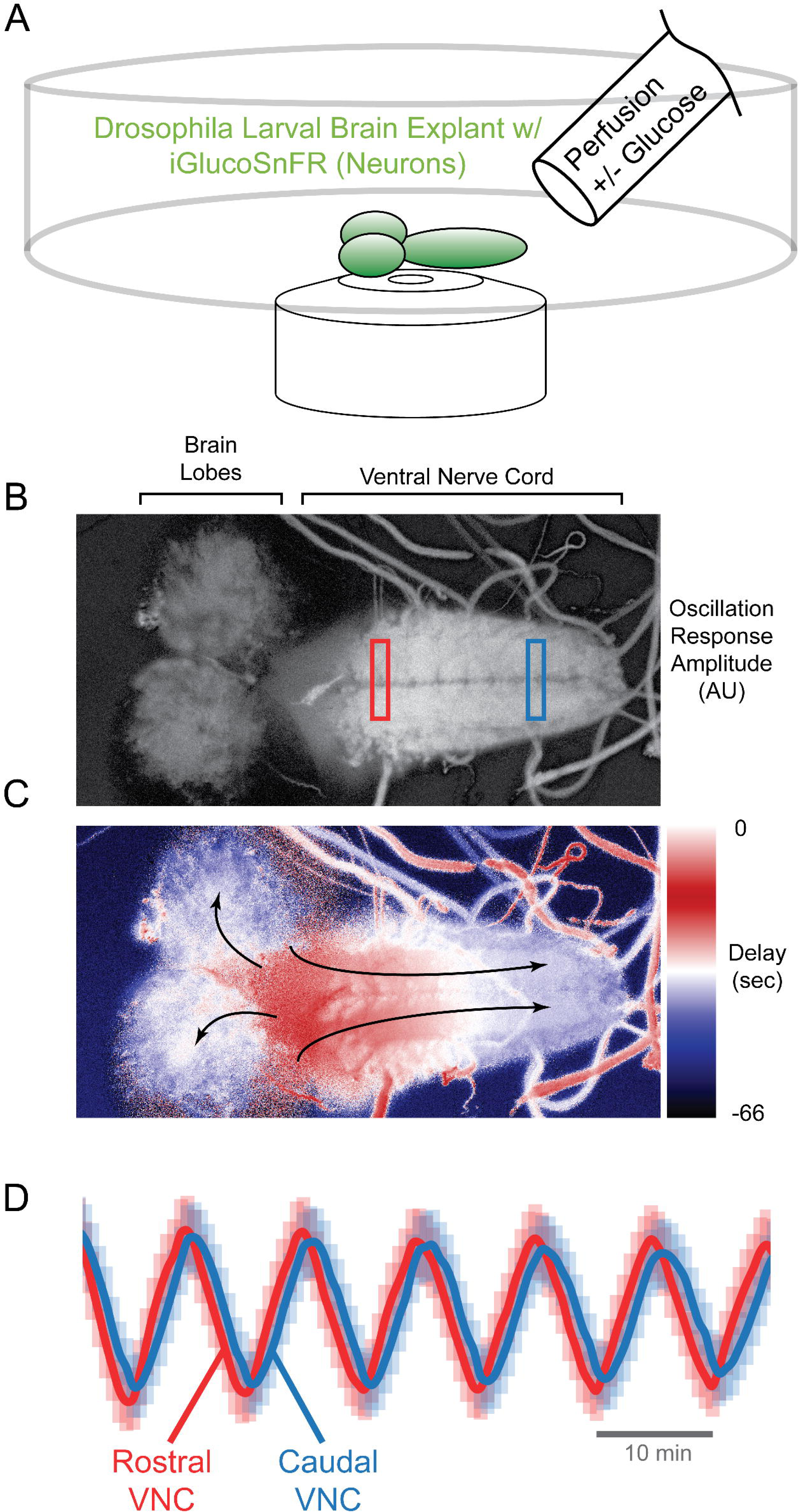
Sensor application in intact larval fruit fly CNS explant. A: Schematic of assay. Third instar larval fly CNS expressing iGlucoSnFR-mRuby2 in all neurons (57C10 driver), embedded in a thin layer of 1% agar, were subjected to continuous perfusion +/-glucose under continuous volumetric confocal imaging. Eight planes at 4.8 µm steps; two channels: Ex 488 nm, Em 505–550; Ex 561, Em 565–650; objective Plan-Apochromat 20x/0.8 M27, imaging rate ∼9 s per volume; 1024 x 512 pixels; stimulus period 10 min. B: FFT amplitudes of response. Time lapse intensity-normalized average-projections in Z were subjected to pixelwise temporal FFT. Magnitudes at stimulus frequency were extracted, and represented in greyscale. C: FFT phases/delays of response. Similar to B, but here stimulus frequency phases were extracted, revealing a temporal gradient in glucose response along the axis of the VNC neuropil and to a lesser extent in the brain lobes (arrows indicate direction of response). D: ΔF/F_avg_ traces from VNC. Normalized fluorescence traces from rectangular regions of interest at rostral or caudal ends of VNC as indicated. Note phase shift. Error bars represent standard deviations of pixels within each ROI.

### In vivo *characterization in zebrafish larvae*

Transparent larval zebrafish are excellent preparations for *in vivo* imaging; furthermore, the ease with which they take up drugs from their water makes simultaneous imaging and pharmacology straightforward. A stable zebrafish transgenic line, Tg (*actb2* :: iGlucoSnFR), was generated to express iGlucoSnFR broadly throughout the animal using the *β*-actin promoter. Larval (5-6 dpf) fish were embedded in agar (no anesthesia or paralysis) and imaged volumetrically on a confocal microscope (15 sections at 15 µm thickness, 512 x 256 pixels, ∼10 seconds/volume). During initial characterization, it became apparent that not only did spontaneous muscle twitches of the agar-mounted fish affect intracellular glucose in muscle, but also that these responses were different in fed versus unfed fish. Thus, a stable Tg (*acta1a* :: jRGECO1a) line was generated as well, expressing the red genetically encoded calcium indicator jRGECO1a^39^ in skeletal muscle with the *α-actinin* promoter. Crossing these two lines yielded Tg (*actb2* :: iGlucoSnFR; *acta1a* :: jRGECO1a), allowing simultaneous measurement of glucose uptake (green) into all cells and action potential firing and accompanying Ca^2+^ uptake (red) into muscles.

Confocal imaging of the double transgenic line revealed intriguing relationships between muscle twitching and glucose uptake in different body regions. In unfed fish, calcium spikes in muscle were always followed by a graded rise in intracellular glucose, which then gradually returned to baseline (**Fig. 4a-top, Supp. Movie 3a**). The largest glucose uptake events seemed to follow the largest muscle spikes (**Fig. 4a-top**). (We did not quantitate this effect due to the relatively slow temporal resolution of the volumetric confocal imaging, which hampered measurement of the exact magnitude of jRGECO1a spikes.) In fed fish, however, glucose levels appeared steady regardless of muscle twitching (small, short-lived responses were seen, but might be movement artifacts; **Fig. 4a-bottom**). This is likely due to higher extra- and intracellular resting glucose levels enabling cells to recover more quickly in the fed state.

**Figure 4:**
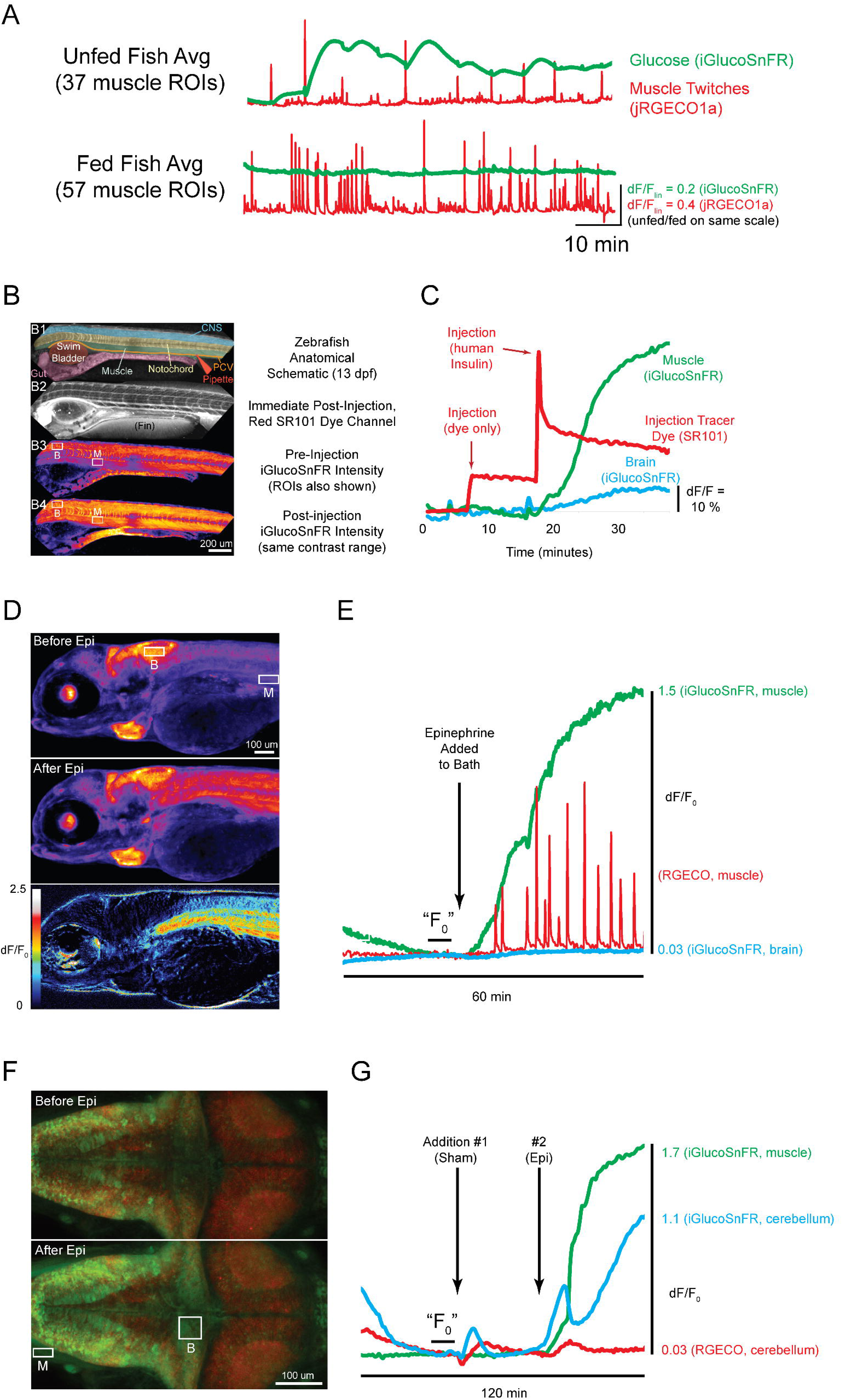
Sensor application in zebrafish muscle and brain *in vivo*. A: Glucose responses to spontaneous muscle twitches. We generated a zebrafish line that expresses iGlucoSnFR broadly (*β*-actin promoter) and the red calcium indicator jRGECO1a in muscle (*α*-actinin promoter). Non-anesthetized, non-paralyzed fish were imaged 5-6 days post fertilization (dpf) for spontaneous responses to muscle twitches. Representative averaged traces from volumetric time-lapse confocal imaging of fed and never-fed fish are presented as indicated. B: Imaging of insulin responses. 13 dpf zebrafish broadly expressing iGlucoSnFR (*β*-actin promoter). B1: schematic illustrating anatomy. Note injection pipette poised for injection into the posterior cardinal vein (PCV). B2: Red fluorescence channel immediately post-injection with insulin and sulforhodamine-101 (SR101) as a marker dye. The entire vasculature became transiently fluorescent red after injection, demonstrating efficacy of systemic injection. B3-4: iGlucoSnFR fluorescence before and after insulin injection. ROIs on brain (“B”) and muscle (“M”) as indicated are plotted in C. (two channels: Ex 488 nm, Em 505–550; Ex 561, Em 565– 650; objective EC Plan-Neofluar 10×/0.3 NA M27, imaging rate ∼1 s per image; 512 x 512 pixels). C: Insulin-induced fluorescence changes. Red trace, injection of dye or dye plus insulin. Green trace, iGlucoSnFR fluorescence measured from muscle (ROI M in panel B). Blue trace, iGlucoSnFR fluorescence measured from brain (ROI B in panel B). Small, artifactual changes in fluorescence appear with introduction of the pipette (dye only injection). Sustained fluorescence increases arise after insulin injection. D: Imaging organism-scale epinephrine responses. 5 dpf fish expressing both iGlucoSnFR (all cells, *β*-actin promoter) and jRGECO1a (muscle cells, *α*-actinin promoter). Top two images (before and after epinephrine) show false color raw intensities from a volumetric time-lapse movie average-projected in Z. ROIs on muscle (“M”) and brain (“B”) are plotted in E. Bottom image shows ΔF/F_0_ after addition of epinephrine, with muscle showing the greatest changes. (5 dpf; fifteen planes at 15 μm steps; two channels: Ex 488 nm, Em 505–550; Ex 561, Em 565–650; objective EC Plan-Neofluar 10×/0.3 NA M27, imaging rate ∼10 s per volume; 512 x 256 pixels). E: Epinephrine-induced fluorescence changes. Blue trace, iGlucoSnFR fluorescence measured from hindbrain (ROI B in panel D). Green trace, iGlucoSnFR fluorescence measured from muscle (ROI M in panel B). Red trace, jRGECO1a fluorescence measured from the same muscle ROI. F_0_ was defined as the averaged intensity values just before addition of epinephrine to the bath, as indicated. Note delayed increase of neuronal activity subsequent to glucose increases. F: Imaging brain-scale epinephrine responses. Single plane (after stack registration) of a 5 dpf fish expressing both iGlucoSnFR (all cells, *β*-actin promoter) and jRGECO1a (neurons, elavl3 promoter) before and after epinephrine addition. ROIs on muscle (“M”) and cerebellum (“B”) are plotted in G. (5 dpf; forty planes at 3.3 µm steps; two channels: Ex 488 nm, Em 505–550; Ex 561, Em 565–650; objective Plan-Apochromat 20x/0.8 M27, imaging rate ∼25 s per volume; 512 x 256 pixels). G: Epinephrine-induced fluorescence changes in the brain. Green trace, iGlucoSnFR fluorescence measured from muscle (ROI “M” in panel F). Blue trace, iGlucoSnFR fluorescence measured from cerebellum (ROI “B” in panel F). Red trace, jRGECO1a fluorescence from same cerebellum ROI. F_0_ was derived from the averaged intensity values just before addition of epinephrine to the bath.

The hormone insulin increases glucose uptake in skeletal muscles. Upon injection of human insulin directly into the posterior caudal vein of Tg (*actb2* :: iGlucoSnFR) fish, large increases in intramuscular glucose were observed (**Fig. 4b-c, Supp. Movie 3b**) with a rise time to half-maximum of about 4-6 minutes (**Fig. 4c**), consistent with results in cultured muscle cells^40^. Saline-only injections did not elicit any response (**Fig. 4c:** *early part of traces*). The hormone epinephrine also has a potent effect on glucose metabolism in muscles: it stimulates glycogenolysis and inhibits glycogen production, leading to high intracellular concentrations of glucose-6-phosphate to feed into augmented glycolysis. At the same time, epinephrine increases systemic glucose rapidly through mobilization of glycogen stores in the liver into glucose followed by efflux into the circulatory system. Addition of epinephrine to the bath of Tg (*actb2* :: iGlucoSnFR; *acta1a* :: jRGECO1a) fish led to dramatic increases in intramuscular glucose levels within about two minutes (**Fig. 4d-e, Supp Movies 3c-e**), presumably the time necessary to diffuse through the agar and into the fish. About five minutes after the initial rise in intramuscular glucose, the fish began to twitch more frequently, and jRGECO1a indicated more muscle spiking (**Fig. 4e**), consistent with “physiological tremor” known to be induced by epinephrine^41^. It was noteworthy that glucose increases began several minutes before calcium spikes appeared, allowing for the possibility that glucose itself might play a role in spawning the observed muscular fasciculations.

In a second set of experiments, epinephrine effects on brain function were monitored by imaging Tg (*actb2* :: iGlucoSnFR; *HuC/elavl3* :: jRGECO1a) fish. Upon a sham addition of fish water into the bath, several brain regions (notably cerebellum and hindbrain; **Fig. 4f**) showed immediate responses in both imaging channels, indicating transient increases in neuronal firing and glucose uptake (**Fig. 4g, Supp. Movies 3f-g**). Muscle tissue in the same field of view showed no response to sham addition. Subsequent addition of epinephrine to the bath led to immediate jRGECO1a and iGlucoSnFR responses in the brain, similar to the sham addition. The fluorescence of brain-expressed iGlucoSnFR showed a larger, prolonged increase, as did muscle-expressed iGlucoSnFR (**Fig. 4g**). Sample deformation in the brain upon epinephrine addition precluded precise mapping of intensity changes, but visual inspection and cross-checking intensity changes of neighboring regions in X, Y, and Z dimensions indicated that the changes were large and not artifactual.

## Discussion

We have presented the design, optimization and deployment of a series of genetically encoded, single-wavelength sensors for glucose (with affinities ranging from 1 µM to 10 mM). They can be easily targeted to specific populations of cells and to sub-cellular compartments to visualize changes in glucose concentration in preparations from cultured cells to intact living animals. They are amenable to long-term expression and imaging. The microbial nature of the binding protein likely renders them otherwise biologically inert in plants and animals. Due to the high glucose concentrations in most situations of interest, buffering by the sensor is negligible. Their single-wavelength nature makes them easy to image and compatible with simultaneous imaging of other physiological phenomena in a separate color channel. Fusion of the red fluorescent protein mRuby2 allows ratiometric imaging for samples with motion artifacts, or simultaneous imaging of two cell types, as shown here with neurons and astrocytes.

The use of iGlucoSnFR with the oscillating stimulus transporter assay (OSTA)^32^ allows precise and rapid measurement of glucose transporter properties. Here we showed both Glut1-mediated glucose transport and inhibition by cytochalasin B, which recapitulated previous determinations of IC_50_. Typically, glucose transporters have been studied with radiolabeled substrates or fluorescent dyes. Radioisotope measurements are hampered by poor spatial and temporal resolution, and thus preclude longitudinal monitoring of transport rates. Fluorescent glucose analogs have different rates of transport between transporters relative to unmodified glucose and can produce misleading results. Robust genetically encoded sensors make these measurements straightforward. The ability of the sensors to produce large fluorescent responses to unmodified glucose ensures accuracy of the results. The amenability of the sensors to prolonged longitudinal imaging also facilitates the tracking of glucose transport across a range of physiological perturbations, such as cytosolic signaling pathways, extracellular factors, pharmacological agents, or endo/exocytosis of transporter populations.

iGlucoSnFR revealed greater glucose uptake in astrocytes than in neurons in co-culture. This phenomenon has been previously described using fluorescent glucose analogs^36,41^, but those results were ambiguous due to the glucose analog having different transport rates among transporter types (*i.e.*, between Glut3, the primary neuronal glucose transporter, and Glut1, the primary astrocyte glucose transporter). The differences in glucose transport rates observed in culture may have ramifications for the ongoing discussion concerning the nature of neuronal metabolism, and particularly the astrocyte-neuron lactate shuttle hypothesis^42^.

*Drosophila* primarily use the glucose disaccharide trehalose (α1-D-glucopyranosyl-α1-D-glucopyranoside) as their hemolymph sugar^43^, but do maintain circulating glucose at low levels. Energy transduction is through glucose, which can be rapidly created through trehalose degradation. In *Drosophila* CNS explants, iGlucoSnFR showed that, contrary to expectations for a simple diffusion model of glucose uptake, interior parts of the CNS saw increases in intracellular glucose concentration before bath-exposed, peripheral regions. It is important to note that these oscillations are distinct from the spontaneous neural activity rhythms generated in explanted larval fly nervous systems^44^. The spontaneous rhythms are relatively rapid and can proceed in either the rostral-to-caudal or caudal-to-rostral direction, depending on forward or reverse fictive crawling. The glucose-induced uptake rhythms, however, proceed in lockstep with the perfused glucose oscillation period (10 min). The delay in brain glucose uptake is consistent with the blood-brain barrier (BBB) being intact in the preparation. It remains to be seen whether the observed rostro-caudal glucose pathway is present *in vivo*, what its mechanism and function might be, and where and how glucose enters the CNS. Since the origin of the apparent glucose waves is near the esophageal foramen, which is surrounded by glucose-relevant DILP neurons^45^, it may be speculated that the esophageal foramen provides a direct port of entry for glucose into the CNS. Identification of spatiotemporal glucose trafficking patterns such as this is currently only possible through the use of iGlucoSnFR and other similar sensors; it will be interesting to see what other spatiotemporal patterns will be found through its use.

In zebrafish larvae, iGlucoSnFR reported insulin- and epinephrine-induced increases in brain and muscle glucose. These changes paralleled neuronal and muscle activity as reported by multiplexed imaging with the calcium sensor jRGECO1a. We expect higher resolution correlations between calcium and glucose signals to be achieved with faster and more spatially focused volumetric imaging^46^. Zebrafish expressing iGlucoSnFR could provide an experimentally tractable method for investigating the details of exercise-mediated glucose uptake in muscle, which is obviously of high medical relevance to both obesity and diabetes.

Many important questions about glucose regulation remain incompletely answered. What are the precise molecular mechanisms by which circulatory glucose levels are correctly sensed, leading to insulin or glucagon secretion from the pancreas? What is the contribution of the various pancreatic cell types^47,48^? What initiates impaired glucose tolerance and how does this result in insulin resistance? What is the role of intestinal gluconeogenesis and how is gut-brain-pancreas glucose sensing coordinated^49^? What is the interplay between glucose dyshomeostasis and autoimmune disease^50^ ? These questions need tools for measuring glucose with high spatial and temporal resolution, and iGlucoSnFR will undoubtedly be useful for their study.

## Methods

### Cloning & mutagenesis

The gene for TtGBP, lacking the periplasmic leader sequence, and codon optimized for expression in *E. coli*, was synthesized by PCR assembly from 42 bp fragments and cloned into pRSET-A (Invitrogen) by BamHI/EcoRI digest (with BamHI encoding Gly-Ser). The gene for cpGFP was inserted, with Gly-Gly linkers on either end, between residues 326 and 327 of TtGBP by overlapping PCR. Single and double amino acid mutations were made by the uracil template-based method^51^.

### Protein characterization

Protein was expressed by overnight growth at 30°C in 0.5 L auto-induction media^52^ and pelleted by centrifugation, resuspended in PBS, and frozen overnight. Cells were lysed by thawing and sonication. Lysate was clarified by centrifugation at 35,000g and purified by immobilized metal affinity chromatography (IMAC) on a 5 mL Fast Flow Chelating Sepharose column (GE Life Sciences) with an elution gradient from 0 to 200 mM imidazole over 120 mL. Clear fractions with absorbance at 280 nm were pooled, concentrated by spin concentrator, and dialyzed 5x into PBS. Non-fluorescent TtGBP (wild-type), and mutants H66A, H348A, and the double mutant H66A+H348A were tested for D-glucose binding affinity by isothermal titration calorimetry. Briefly, 20 µM purified protein was titrated with a concentrated stock of D-glucose in a MicroCal ITC. Wild type TtGBP showed an affinity of about 1 µM, and the mutants were progressively weaker.

### Mutant screening

TtGBP(H66A, H348A)-cpGFP was used as the template for mutagenesis. Mutations were targeted to the Gly-Gly linkers between of TtGBP and cpGFP. Mutated plasmids were transformed into T7 Express cells (New England BioLabs), plated on LB+100 µg/mL ampicillin agar plates and grown overnight at 37°C. Colonies were picked into 2 mL 96-well plates containing 0.9 mL auto-induction media^52^ with 100 µg/mL ampicillin and grown overnight with vigorous shaking (400 RPM) at 30°C. Bacterial pellets were collected by centrifugation (3000g), and resuspended in 0.5 mL PBS by vortexing to wash away endogenous glucose, and pelleted again. The wash procedure was repeated three times, and pellets were frozen overnight at −20°C. Frozen bacterial pellets were then lysed by addition of 1 mL PBS and vortexing. Cellular debris was pelleted by centrifugation and clarified lysate removed for fluorescence assays by pipetting.

The green fluorescence (Ex 485 nm, Em 515 nm, 5 nm bandpass) of 100 µL of clarified lysate was measured in a Tecan Safire 2 plate-reading fluorimeter. D-Glucose was added to a final concentration of 10 mM and fluorescence measured again. Variants with increased ΔF/F over the starting construct were immediately re-assayed to estimate binding affinity, and winners were streaked out on agar plates, re-isolated, re-assayed, and sequenced. After screening of about 800 variants at each linker, the variant with Linker 1 – PA, Linker 2 – NP was chosen as iGlucoSnFR because it had the highest ΔF/F.

### Cultured HEK293 cells

Cell culture: Glut1 experiments were done on HEK293 (ATCC #CRL-1573) cells transfected using the Amaxa system (Lonza) and 1 µg DNA. Cells were cultured after transfection to densities of 50%–90% confluence in 35 mm coverslip-bottomed culture dishes (MatTek).

### Cultured neurons/astrocytes

Hippocampi were dissected from P0 neonatal wild-type Sprague-Dawley rat pups (Charles River) in dissection buffer (500 ml HBSS, 10 mM HEPES, 100 U/mL Penicillin and 10 µg/mL Streptomycin (GIBCO 15140-122)), cut into small pieces and dissociated in papain enzyme (Worthington, #PAP2) in dissection buffer for 30 minutes at 37°C. After 30 min of enzyme digestion, papain enzyme solution was removed and tissue pieces dissociated by trituration in MEM (Invitrogen GIBCO 51200-038, no phenol red) with 10% FBS media (500 ml of MEM, 2.5 g D-glucose, 100 mg NaHCO_3_, 50 mg transferrin (Calbiochem 616420), 50 ml FBS (heat inactivated; HyClone SH30071.03 HI), 5 ml 0.2 M L-glutamine solution, 12.5 mg human insulin (Sigma I-6634), 100 U/mL Penicillin and 10 µg/mL Streptomycin (GIBCO 15140-122)). Resulting cell suspension was filtered through a 70 µm filter to remove remaining pieces of tissue and centrifuged at 90 x g for 7 minutes to pellet. The cell pellet was re-suspended in plating media (500 ml of MEM, 2.5 g D-glucose, 100 mg NaHCO_3_, 50 mg Transferrin, 50 ml FBS, 5 ml 0.2 M L-glutamine solution, 12.5 mg insulin, 100 U/mL Penicillin and 10 µg/mL Streptomycin) and counted for cell viability and counts. Cells were plated at 35,000 cells/coverslip with 70 µl plating media in center of a coverslip-bottomed cultured dish (Mattek) and incubated at 37°C for attachment to poly-D-lysine-coated coverslips. After 2+ hours (once cells attached), 1 ml of NbActiv4 media (BrainBits) was added and cultured for two weeks. Media changes were done by removing 0.5 ml and adding 0.5 ml fresh NbActiv4 media. Resulting cultures were infected with AAV viruses one week after plating, and were further cultured for 2-3 weeks prior to experiments.

### Perfusion system

A gravity-fed, four-channel perfusion system (VC3-4, ALA Scientific Instruments) was used in all perfusion experiments. The bottoms of the 60 ml feeder syringes were raised to a height which provided flow rates of 4–6 mL/min depending on the fluid level of the syringes. The outlet of the system was directed toward the illuminated area of the coverslip dish at a distance of 1–3 mm, and continuous fast suction at the raised edge of the dish removed solutions. The perfusion outlet was Tygon tubing with 1/16’’ inner diameter (somewhat larger than that in the manifold), which might have slowed the velocity of the buffers, allowing for less disturbance at the sample. Protocols for buffer switching were carried out by the control software provided. Buffer changes were characterized to be > 90% in 1 second.

### AAVs

Adeno-associated viruses were generated of serotype 2/1 using promoters for neurons (*hSynapsin1*) or glia (*gfABC*_*1*_*D*), and containing either the cytosolic sensor alone (iGlucoSnFR) or the mRuby2-tagged cytosolic sensor (iGlucoSnFR-mRuby2). Viral titers were 1-2 x 10^13^ GC/mL.

### Flies

Virgin female Drosophila melanogaster carrying the intracellularly-targeted iGlucoSnFR transgene (LexAop-iGlucoSnFR; +; +) were crossed with males from a driver line (w/y; 57C10-LexA; +)^38^ that expresses pan-neuronally in post-embryonic flies. Larvae were raised on normal cornmeal-molasses fly food. Third instar larvae were manually selected and the CNS was dissected in physiological saline^46^. The CNS explants were placed in a 35 mm coverslip-bottom culture dish. To hold the samples stationary, they were partially embedded in a thin layer of 1% low-melting point agarose (SeaPlaque, Lonza) dissolved in physiological saline. After the agarose cooled and polymerized, the embedded CNS explant was then submerged in physiological saline.

### Fish

Fish were generated using Tol2 transgenesis^53^. Transgenic fish Tg(*actb2* :: iGlucoSnFR) express iGlucoSnFR under the *β*-actin promoter, and hence target nearly all cells. Tg(*acta1a* :: jRGECO1a) express the red calcium indicator jRGECO1a in skeletal muscle cells^54^. For insulin injection experiments, a non-functional electrophysiological glass patch pipette was mounted on a micromanipulator, and was filled with insulin or blank solutions, along with 1 µM of the red fluorescent dye sulforhodamine 101. This pipette was positioned outside the fish near to the posterior caudal vein prior to the beginning of imaging and a baseline of 5-10 minutes of imaging was collected. Imaging was then paused briefly (10-30 seconds) while the pipette was inserted. Imaging was immediately restarted, and dye was injected by manual pressure, verified by presence of bright red fluorescence, then the pipette was retracted. When necessary, this pipette was changed to a new pipette containing a different solution, and the procedure repeated in the same imaging session.

### Imaging and Image Processing: HEK cells

Images of HEK cells co-expressing hGlut1 (Addgene #18085) and iGlucoSnFR.mRuby2 were acquired on an inverted Zeiss LSM 800 confocal microscope equipped with GaAsP detectors and an EC Plan-Neofluar 10x/0.30 M27 objective. Images were 512 x 512 pixels at 16-bit depth, with a frame time of 933 ms. Excitation used laser lines at 488 and 561 nm, with emission ranges of 495-575 and 575-617 nm. The red channel served to confirm that cells were stable during the experiment in the x-y and focal planes, but was omitted from further processing. To separate the oscillating component of the signal from gradual baseline drifts or trends, the green (sensor) channel was processed by subtracting a two-sided moving-window average (F_avg_) corresponding to the period length of the oscillating stimulus, a common de-trending technique. Regions of interest (ROIs) corresponding to co-transfected cells were selected by thresholding an image composed of the oscillation magnitudes at each pixel during the uniform pre-treatment epoch of the experiment, and signals from the detrended data were extracted in these ROIs. Individual ROI traces were then normalized by division by the RMSD of the uniform initial epoch of the experiment, and are presented in plots as individual data points. For fitting IC_50_, percent inhibition was calculated as 100*[1 - RMSD_before_/RMSD_after_] with equal length windows for RMSD calculations.

### Imaging and Image Processing: Neuron/astrocyte co-cultures

Images of hippocampal cultures infected with both AAV2/1-*hSynapsin1*-iGlucoSnFR-mRuby2 and AAV2/1-*gfaABC*_*1*_*D*-iGlucoSnFR or AAV2/1-*hSynapsin1*-iGlucoSnFR and AAV2/1-*gfaABC*_*1*_*D*-iGlucoSnFR-mRuby2 were acquired on an inverted Zeiss LSM 800 confocal microscope equipped with GaAsP detectors and an EC Plan-Neofluar 20x/0.80 M27 objective. Images were 1024 x 512 pixels at 16-bit depth, with a frame time of 933 ms. Excitation used laser lines at 488 and 561 nm, with emission ranges of 495-575 and 575-617 nm. The red channel served to differentiate cell types. Images were processed by first average-binning over time by a factor of two, followed by detrending to separate the oscillating component (ΔF) of the signal from gradual baseline drifts or trends (F_avg_). The experimental green (sensor) channel was processed by subtracting F_avg_ from the original image stack, yielding ΔF, then divided by F_avg,_, yielding a ΔF/F_avg_ image stack. Regions of interest (ROIs) over mRuby2-expressing cells were selected by thresholding the red channel F_avg_ image stack using a reasonably low (mean) threshold, converting to a binary mask, then minimum-projected to identify regions that were consistently fluorescent in the red channel over the course of the experiment. The green F_avg_ stack was then multiplied by this mask and minimum-projected, then thresholded by maximum entropy to identify remaining fluorescent green ROIs. ROIs greater than 50 pixels in size were identified, and mean intensities under these ROIs were evaluated in the green channel ΔF/F_avg_ image stack. These traces were then averaged and presented. The somewhat complex nature of this processing procedure was designed in the hopes of identifying and differentiating with highest certainty cells expressing iGlucoSnFR and not iGlucoSnFR-mRuby2, and were consistently fluorescent over the course of the experiment (a few cells disappeared, detached, or moved).

To identify cells that expressed iGlucoSnFR-mRuby2 and not iGlucoSnFR, the F_avg_ stack in the red channel was thresholded using the “moments” threshold in FIJI/ImageJ, followed by minimum projection and identification of ROIs which were consistent fluorescent in the red channel. These ROIs were then evaluated in green channel ΔF/F_avg_ stack and presented.

The exact same processing procedure was used in both types of experiments: AAV2/1-*hSynapsin1*-iGlucoSnFR-mRuby2 with AAV2/1-*gfaABC*_*1*_*D*-iGlucoSnFR or AAV2/1-*hSynapsin1*-iGlucoSnFR and AAV2/1-*gfaABC*_*1*_*D*-iGlucoSnFR-mRuby2.

Raw, time-averaged, mean intensity values in the green channel of all ROIs were measured as well, and no obvious correlation of intensity with oscillation magnitudes was observed.

### Imaging and Image Processing: Drosophila CNS

Images of explanted 3^rd^ instar CNSes expressing iGlucoSnFR.mRuby2 pan-neuronally (57C10 driver) were acquired on an inverted Zeiss LSM 800 confocal microscope equipped with GaAsP detectors and an EC Plan-Neofluar 20x/0.8 M27 objective. Data were collected as eight x 5 µm planes at 1024 x 512 pixels at 16-bit depth, with a volume rate of ∼9 s. Excitation used laser lines at 488 and 561 nm, with emission ranges of 505-550 and 565-650 nm. The red channel, which remained roughly stable in intensity over the course of the experiment, served to confirm the glucose specificity of signal as well as allow for ratiometric imaging, which was found to be unnecessary. The same moving-window subtraction detrending process described above was used to accentuate oscillations, and drew attention to the rostro-caudal glucose increase pattern described. Pixelwise temporal FFT analysis allowed plotting of the graded phase shifts along the rostro-caudal axis.

### Imaging and Image Processing: Zebrafish Larvae

For data on muscle, imaging was carried out as described above. To extract signals in the spontaneous twitch experiments, image stacks were first registered over time by the green channel. ROIs were then identified by first normalizing local contrast in the image stack and thresholding. These ROIs were tracked over time in the original raw image stacks, and mean fluorescence intensities in each ROI in both green and red channels were quantified, then averaged together and plotted. For both insulin and epinephrine data, images were first registered in the green channel, then a simple rectangular ROI was selected for simplicity of presentation. In the epinephrine experiments, there was some deformation of the subject upon addition of epinephrine, complicating presentation of a “clean” still image displaying fluorescence changes. A rectangular ROI was selected based on relative structural stability of the region as well as fluorescence intensity changes. Neighboring regions in XYZ were confirmed not to manifest opposing/mirrored responses (which would suggest that the observed signal was a motion artifact).

## Supporting information

Supp Mov 1a

Supp Mov 1b

Supp Mov 2a

Supp Mov 2b

Supp Mov 2c

Supp Mov 2d

Supp Mov 3b

Supp Mov 3c

Supp Mov 3f

Supp Mov 3g

## Acknowledgements

We would like to thank Jay Unruh for ImageJ plugins; Gary Yellen, Jim Truman, Tim Ryan, and Phil Borden for helpful discussions; Deepika Walpita for cell culture; Janelia Fly Facility for fly maintenance; and Jared Rouchard & the Janelia Aquarium Facility for fish maintenance.

## Author contributions

JPK, JSM, SK, RTL, and LLL conceived, designed, and constructed the sensor and variations thereof. JPK, JSM, and SK characterized the sensor in purified protein. JPK, JSM, HL, WCL, JS, MK, PJK, and LLL contributed to the design of cellular & animal experiments. JPK, JSM, HL, and WCL generated experimental animals, prepared, and/or conducted physiological experiments. JPK, JSM, WCL, MK, and LLL analyzed the data. JPK, JSM, and LLL wrote the manuscript.

## Code availability

All analysis code used in this study is available upon request.

## Data availability

All data from this study is available upon request.

## Reagent availability

DNA constructs are available from Addgene. Transgenic fly lines have been deposited at Bloomington (**will be updated**). Transgenic fish lines are available upon request.

## Accession codes

All constructs have been deposited at Addgene (**will be updated**). Sequences have been deposited in Genbank (**will be updated**).

