## Supplementary Information for "*In vivo* glucose imaging in multiple model organisms with an engineered single-wavelength sensor"

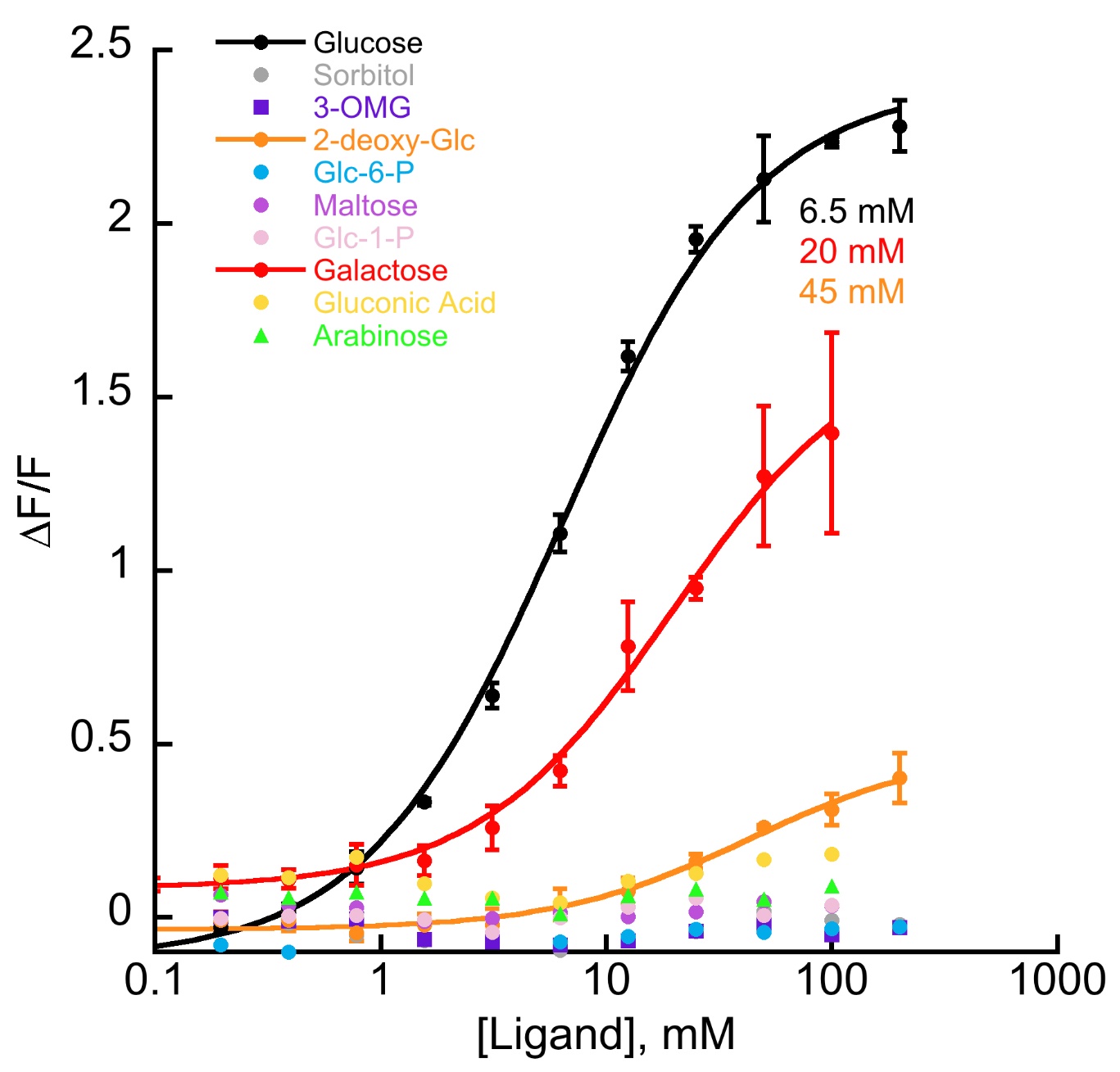

**Supplementary Figure 1. Titrations of glucose-related ligands**. Glucose bound most tightly, with a K_d_ of 6.5 +/- 0.4 mM, and only galactose and 2-deoxy-glucose showed any fluorescence changes (K_d_’s = 20 +/- 3 and 45 +/- 7 mM, respectively). 3-OMG is 3-O-methyl-glucose, Glc-6-P is glucose-6-phosphate, Glc-1-P is glucose-1-phosphate. Traces are mean of n=3 experimental replicates; error bars are std. dev.

**Supplementary Figure 2. iGlucoSnFR pH dependence.** Sensor was titrated over a range of pH values, revealing a pH dependence both of K_d_ and ΔF/F, with larger pH values tending to increase both. Traces are mean of n=4 experimental replicates; error bars are std. dev.
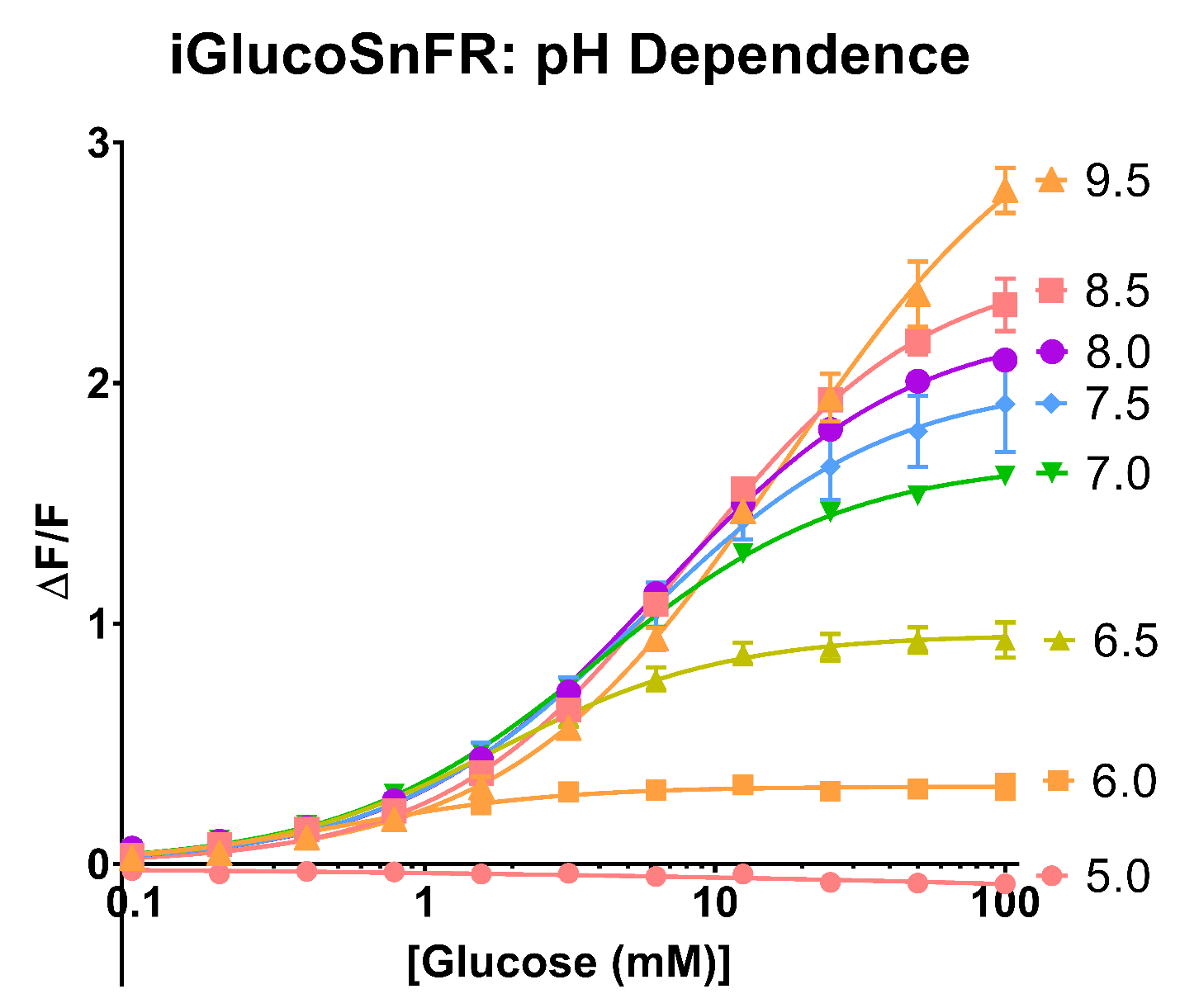

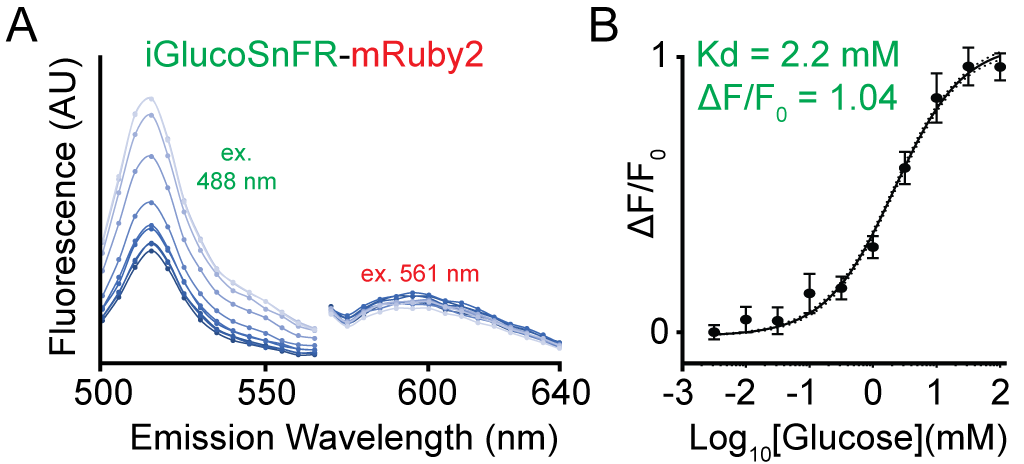

**Supplementary Figure 3. *In vitro* characterization of iGlucoSnFR-mRuby2**. iGlucoSnFR tagged with mRuby2 in the same polypeptide was expressed in *E. coli* and purified. A. Emission spectra in the green emission band at various glucose concentrations were indistinguishable from the green-only sensor. Red emission spectra were unresponsive to glucose and were indistinguishable from mRuby2. B. iGlucoSnFR responses to glucose in the green emission band were somewhat different from those in the green-only sensor. As mentioned in the main text, this construct was difficult to purify to homogeneity when expressed bacterially, so it is hard to be confident in the accuracy of these titrations, especially in light of the cellular data showing identical responses to identical stimuli. Overall n = 60: 3 replicate titrations were measured 4 times each at 5 wavelengths on the green peak. ΔF/F for each titration, measurement, and wavelength, were calculated first, then fit by the Hill equations as 60 titrations. Error bars represent mean and standard deviation.

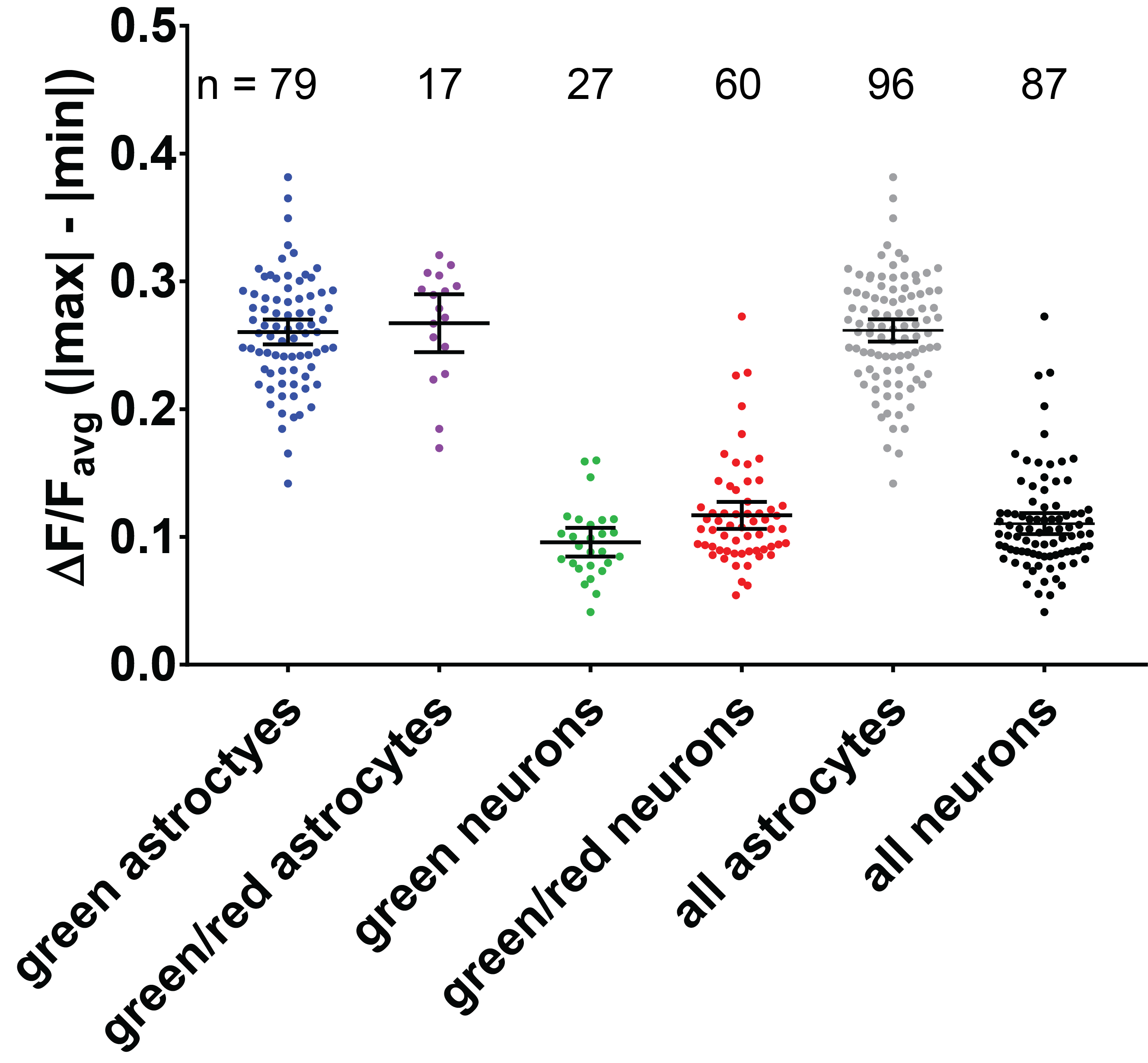

**Supplementary Figure 4. Astrocyte/neuron experiments with iGlucoSnFR and/or iGlucoSnFR-mRuby2**. Astrocyte/neuron co-cultures were infected with either the combination (AAV2/1-*hSynapsin1*-iGlucoSnFR and AAV2/1- *gfaABC_1_D*-iGlucoSnFR-mRuby2), which yielded green neurons and green/red astrocytes, or (AAV2/1-*hSynapsin1*-iGlucoSnFR-mRuby2 and AAV2/1- *gfaABC_1_D*-iGlucoSnFR), which yielded green/red neurons and green astrocytes. Responses were imaged and quantified identically in both cases, and showed that the presence or absence of the tag did not change the observed magnitudes of oscillation. The difference between all-astrocytes and all-neurons was significant (p < 10^-16^, Mann-Whitney test). All data points (corresponding to cells/ROIs) are shown, along with mean and 95% confidence intervals.

**>cyto-iGlucoSnFR**

**MGDRSKLEIFSWWAGDEGPALEALIRLYKQKYPGVEVINATVTGGAGVNARAVLKTRMLGGDPPDTFQVHAGMELIGTWVVANRMEDLSALFRQEGWLQAFPKGLIDLISYKGGIWSVPVNIHRSNVMWYLPAKLKEWGVNPPRTWDEFLATCQTLKQKGLEAPLALGENWTQQHLWESVALAVLGPDDWNNLWNGKLKFTDPKAVRAWEVFGRVLDCANKDAAGLSWQQAVDRVVQGKAAFNVMGDWAAGYMTTTLKLKPGTDFAWAPSPGTQGVFMMLSDSFGLPKGAKNRQNAINWLRLVGSKEGQDTFNPLKGSIAARLDSDPSKYPASHNVYIMADKQKNGIKANFKIRHNIEDGGVQLAYHYQQNTPIGDGPVLLPDNHYLSTQSKLSKDPNEKRDHMVLLEFVTAAGITLGMDELYKGGTGGSMVSKGEELFTGVVPILVELDGDVNGHKFSVSGEGEGDATYGKLTLKFICTTGKLPVPWPTLVTTLTYGVQCFSRYPDHMKQHDFFKSAMPEGYIQERTIFFKDDGNYKTRAEVKFEGDTLVNRIELKGIDFKEDGNILGHKLEYNFNNPNAYGQSAMRDWRSNRIVGSLVHGAVAPESFMSQFGTVMEIFLQTRNPQAAANAAQAIADQVGLGRLGQLQVDEQKLISEEDLNAVGQDTQERFYCENEVGS.**

**TtGBP, Linker, cpGFP, Myc tag, ER export motif**

**>cyto-iGlucoSnFR-mRuby2**

**MGDRSKLEIFSWWAGDEGPALEALIRLYKQKYPGVEVINATVTGGAGVNARAVLKTRMLGGDPPDTFQVHAGMELIGTWVVANRMEDLSALFRQEGWLQAFPKGLIDLISYKGGIWSVPVNIHRSNVMWYLPAKLKEWGVNPPRTWDEFLATCQTLKQKGLEAPLALGENWTQQHLWESVALAVLGPDDWNNLWNGKLKFTDPKAVRAWEVFGRVLDCANKDAAGLSWQQAVDRVVQGKAAFNVMGDWAAGYMTTTLKLKPGTDFAWAPSPGTQGVFMMLSDSFGLPKGAKNRQNAINWLRLVGSKEGQDTFNPLKGSIAARLDSDPSKYPASHNVYIMADKQKNGIKANFKIRHNIEDGGVQLAYHYQQNTPIGDGPVLLPDNHYLSTQSKLSKDPNEKRDHMVLLEFVTAAGITLGMDELYKGGTGGSMVSKGEELFTGVVPILVELDGDVNGHKFSVSGEGEGDATYGKLTLKFICTTGKLPVPWPTLVTTLTYGVQCFSRYPDHMKQHDFFKSAMPEGYIQERTIFFKDDGNYKTRAEVKFEGDTLVNRIELKGIDFKEDGNILGHKLEYNFNNPNAYGQSAMRDWRSNRIVGSLVHGAVAPESFMSQFGTVMEIFLQTRNPQAAANAAQAIADQVGLGRLGQLQVDEQKLISEEDLNAVGQDTQERMVSKGEELIKENMRMKVVMEGSVNGHQFKCTGEGEGNPYMGTQTMRIKVIEGGPLPFAFDILATSFMYGSRTFIKYPKGIPDFFKQSFPEGFTWERVTRYEDGGVVTVMQDTSLEDGCLVYHVQVRGVNFPSNGPVMQKKTKGWEPNTEMMYPADGGLRGYTHMALKVDGGGHLSCSFVTTYRSKKTVGNIKMPGIHAVDHRLERLEESDNEMFVVQREHAVAKFAGLGGGMDELYKFYCENEVGS.**

**TtGBP, Linker, cpGFP, Myc tag, mRuby2, ER export motif**

**>extracellular-iGlucoSnFR**

**METDTLLLWVLLLWVPGSTGDRSKLEIFSWWAGDEGPALEALIRLYKQKYPGVEVINATVTGGAGVNARAVLKTRMLGGDPPDTFQVHAGMELIGTWVVANRMEDLSALFRQEGWLQAFPKGLIDLISYKGGIWSVPVNIHRSNVMWYLPAKLKEWGVNPPRTWDEFLATCQTLKQKGLEAPLALGENWTQQHLWESVALAVLGPDDWNNLWNGKLKFTDPKAVRAWEVFGRVLDCANKDAAGLSWQQAVDRVVQGKAAFNVMGDWAAGYMTTTLKLKPGTDFAWAPSPGTQGVFMMLSDSFGLPKGAKNRQNAINWLRLVGSKEGQDTFNPLKGSIAARLDSDPSKYPASHNVYIMADKQKNGIKANFKIRHNIEDGGVQLAYHYQQNTPIGDGPVLLPDNHYLSTQSKLSKDPNEKRDHMVLLEFVTAAGITLGMDELYKGGTGGSMVSKGEELFTGVVPILVELDGDVNGHKFSVSGEGEGDATYGKLTLKFICTTGKLPVPWPTLVTTLTYGVQCFSRYPDHMKQHDFFKSAMPEGYIQERTIFFKDDGNYKTRAEVKFEGDTLVNRIELKGIDFKEDGNILGHKLEYNFNNPNAYGQSAMRDWRSNRIVGSLVHGAVAPESFMSQFGTVMEIFLQTRNPQAAANAAQAIADQVGLGRLGQLQVDEQKLISEEDLNAVGQDTQEVIVVPHSLPFKVVVISAILALVVLTIISLIILIMLWQKKPRFYCENEVGS.**

**Signal Peptide, TtGBP, Linker, cpGFP, Myc tag, TM helix, mRuby2, ER export motif**

**>extracellular-iGlucoSnFR-mRuby2**

**METDTLLLWVLLLWVPGSTGDRSKLEIFSWWAGDEGPALEALIRLYKQKYPGVEVINATVTGGAGVNARAVLKTRMLGGDPPDTFQVHAGMELIGTWVVANRMEDLSALFRQEGWLQAFPKGLIDLISYKGGIWSVPVNIHRSNVMWYLPAKLKEWGVNPPRTWDEFLATCQTLKQKGLEAPLALGENWTQQHLWESVALAVLGPDDWNNLWNGKLKFTDPKAVRAWEVFGRVLDCANKDAAGLSWQQAVDRVVQGKAAFNVMGDWAAGYMTTTLKLKPGTDFAWAPSPGTQGVFMMLSDSFGLPKGAKNRQNAINWLRLVGSKEGQDTFNPLKGSIAARLDSDPSKYPASHNVYIMADKQKNGIKANFKIRHNIEDGGVQLAYHYQQNTPIGDGPVLLPDNHYLSTQSKLSKDPNEKRDHMVLLEFVTAAGITLGMDELYKGGTGGSMVSKGEELFTGVVPILVELDGDVNGHKFSVSGEGEGDATYGKLTLKFICTTGKLPVPWPTLVTTLTYGVQCFSRYPDHMKQHDFFKSAMPEGYIQERTIFFKDDGNYKTRAEVKFEGDTLVNRIELKGIDFKEDGNILGHKLEYNFNNPNAYGQSAMRDWRSNRIVGSLVHGAVAPESFMSQFGTVMEIFLQTRNPQAAANAAQAIADQVGLGRLGQLQVDEQKLISEEDLNAVGQDTQEVIVVPHSLPFKVVVISAILALVVLTIISLIILIMLWQKKPRMVSKGEELIKENMRMKVVMEGSVNGHQFKCTGEGEGNPYMGTQTMRIKVIEGGPLPFAFDILATSFMYGSRTFIKYPKGIPDFFKQSFPEGFTWERVTRYEDGGVVTVMQDTSLEDGCLVYHVQVRGVNFPSNGPVMQKKTKGWEPNTEMMYPADGGLRGYTHMALKVDGGGHLSCSFVTTYRSKKTVGNIKMPGIHAVDHRLERLEESDNEMFVVQREHAVAKFAGLGGGMDELYKFYCENEVGS.**

**Signal Peptide, TtGBP, Linker, cpGFP, Myc tag, TM helix, mRuby2, ER export motif**

**>secreted-iGlucoSnFR**

**METDTLLLWVLLLWVPGSTGDRSKLEIFSWWAGDEGPALEALIRLYKQKYPGVEVINATVTGGAGVNARAVLKTRMLGGDPPDTFQVHAGMELIGTWVVANRMEDLSALFRQEGWLQAFPKGLIDLISYKGGIWSVPVNIHRSNVMWYLPAKLKEWGVNPPRTWDEFLATCQTLKQKGLEAPLALGENWTQQHLWESVALAVLGPDDWNNLWNGKLKFTDPKAVRAWEVFGRVLDCANKDAAGLSWQQAVDRVVQGKAAFNVMGDWAAGYMTTTLKLKPGTDFAWAPSPGTQGVFMMLSDSFGLPKGAKNRQNAINWLRLVGSKEGQDTFNPLKGSIAARLDSDPSKYPASHNVYIMADKQKNGIKANFKIRHNIEDGGVQLAYHYQQNTPIGDGPVLLPDNHYLSTQSKLSKDPNEKRDHMVLLEFVTAAGITLGMDELYKGGTGGSMVSKGEELFTGVVPILVELDGDVNGHKFSVSGEGEGDATYGKLTLKFICTTGKLPVPWPTLVTTLTYGVQCFSRYPDHMKQHDFFKSAMPEGYIQERTIFFKDDGNYKTRAEVKFEGDTLVNRIELKGIDFKEDGNILGHKLEYNFNNPNAYGQSAMRDWRSNRIVGSLVHGAVAPESFMSQFGTVMEIFLQTRNPQAAANAAQAIADQVGLGRLGQLQVDEQKLISEEDLNAVGQDTQERFYCENEVGS.**

**Signal Peptide, TtGBP, Linker, cpGFP, Myc tag, ER export motif**

**>secreted-iGlucoSnFR-mRuby2**

**METDTLLLWVLLLWVPGSTGDRSKLEIFSWWAGDEGPALEALIRLYKQKYPGVEVINATVTGGAGVNARAVLKTRMLGGDPPDTFQVHAGMELIGTWVVANRMEDLSALFRQEGWLQAFPKGLIDLISYKGGIWSVPVNIHRSNVMWYLPAKLKEWGVNPPRTWDEFLATCQTLKQKGLEAPLALGENWTQQHLWESVALAVLGPDDWNNLWNGKLKFTDPKAVRAWEVFGRVLDCANKDAAGLSWQQAVDRVVQGKAAFNVMGDWAAGYMTTTLKLKPGTDFAWAPSPGTQGVFMMLSDSFGLPKGAKNRQNAINWLRLVGSKEGQDTFNPLKGSIAARLDSDPSKYPASHNVYIMADKQKNGIKANFKIRHNIEDGGVQLAYHYQQNTPIGDGPVLLPDNHYLSTQSKLSKDPNEKRDHMVLLEFVTAAGITLGMDELYKGGTGGSMVSKGEELFTGVVPILVELDGDVNGHKFSVSGEGEGDATYGKLTLKFICTTGKLPVPWPTLVTTLTYGVQCFSRYPDHMKQHDFFKSAMPEGYIQERTIFFKDDGNYKTRAEVKFEGDTLVNRIELKGIDFKEDGNILGHKLEYNFNNPNAYGQSAMRDWRSNRIVGSLVHGAVAPESFMSQFGTVMEIFLQTRNPQAAANAAQAIADQVGLGRLGQLQVDEQKLISEEDLNAVGQDTQERMVSKGEELIKENMRMKVVMEGSVNGHQFKCTGEGEGNPYMGTQTMRIKVIEGGPLPFAFDILATSFMYGSRTFIKYPKGIPDFFKQSFPEGFTWERVTRYEDGGVVTVMQDTSLEDGCLVYHVQVRGVNFPSNGPVMQKKTKGWEPNTEMMYPADGGLRGYTHMALKVDGGGHLSCSFVTTYRSKKTVGNIKMPGIHAVDHRLERLEESDNEMFVVQREHAVAKFAGLGGGMDELYKFYCENEVGS.**

**Signal Peptide, TtGBP, Linker, cpGFP, Myc tag, mRuby2, ER export motif**

**Supplementary Figure 5. Sequences of sensors, colored by motifs as indicated**.

**Appendix: semi-quantitative disentanglement of glucose transport and metabolism *via* OSTA waveform analysis**

Calibration of intracellular biosensors *in situ* remains challenging. For one thing, whatever means are used to control cytosolic conditions (*e.g.,* permeabilization with detergents, porins, or ionophores) will inherently perturb the physiological state in multiple ways. Using glucose transport and metabolism in neuron-astrocyte co-cultures, we here outline a “semi-quantitative” approach to the problem.

In attempting to measure glucose transport in neuron-astrocyte co-cultures or other living cells, however, a further experimental complication is layered on top of the calibration issue. Metabolic consumption of glucose becomes intertwined with transport phenomena, leading to underestimates of uptake and overestimates of efflux. Since the true metabolic consumption rate of each cell is not known, it is essentially impossible to determine the true underlying transport rates, even with ideal calibration. What can be determined trivially and unambiguously, however, are the maximum and minimum influx rates given the observed rates of fluorescence, which will give an idea of the relative transport rates of cellular sub-populations. This, namely putting upper and lower bounds on the transport rates, is what is meant by “semi-quantitative” above.

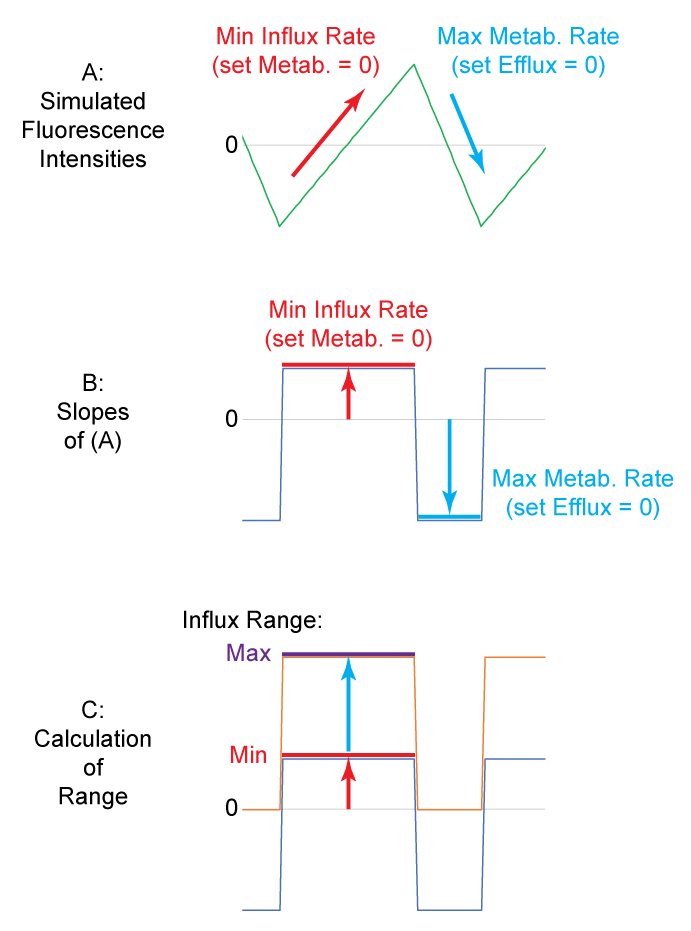
There are a few properties of astrocytes and neurons that should be noted in assessing glucose dynamics. First, glucose transporters in astrocytes (Glut1) and neurons (Glut3) are passive uniporters whose properties are well known: as passive transporters, their transport rates will be governed only by the glucose gradients across them, and concentrative effects are not possible. Therefore, intracellular glucose concentrations will not exceed extracellular concentrations. Second, under given conditions, rates of glucose metabolism will be governed by the concentration of intracellular glucose according to enzyme kinetics, regardless of whether concentrations are rising or falling. Applying these assumptions to OSTA-type data, it can be inferred that the upper bound of metabolic rate is represented by the downward slope in the downstrokes of the assay (**Appendix Figure 1A**, blue arrows). In reality, this downward slope includes a transporter-mediated efflux component as well, but the limiting case would be when none of this negative slope is attributable to efflux. Since this upper bound for metabolic rate is now known, it can be extrapolated to the upstrokes, since each glucose concentration in the downstroke has an analogous point in the upstroke at which the metabolic rates will be equal. In the case of linear or near-linear “sawtooth” waveforms as in the current data, this calculation simplifies to subtracting the constant negative slope (maximum metabolic rate) from the slopes across the entire waveform, yielding a maximum rate for influx in the upstrokes. In this way, under given conditions, a maximum bound for influx rate can also be inferred.

Appendix Figure 1: Theory

The lower bound for influx rate can be also be derived by supposing the metabolic rate to be zero, and attributing the measured fluorescence increase in the upstroke to transport, undiminished by metabolic consumption. This lower bound is therefore simply the measured slope in the upstrokes (**Appendix Figure 1A**, red arrows). These lower and upper bounds define a range of possible influx rates given the observed intensity changes, and when the ranges of two cell populations in a given field of view are compared, a semi-quantitative idea of their relative transport rates can be achieved.

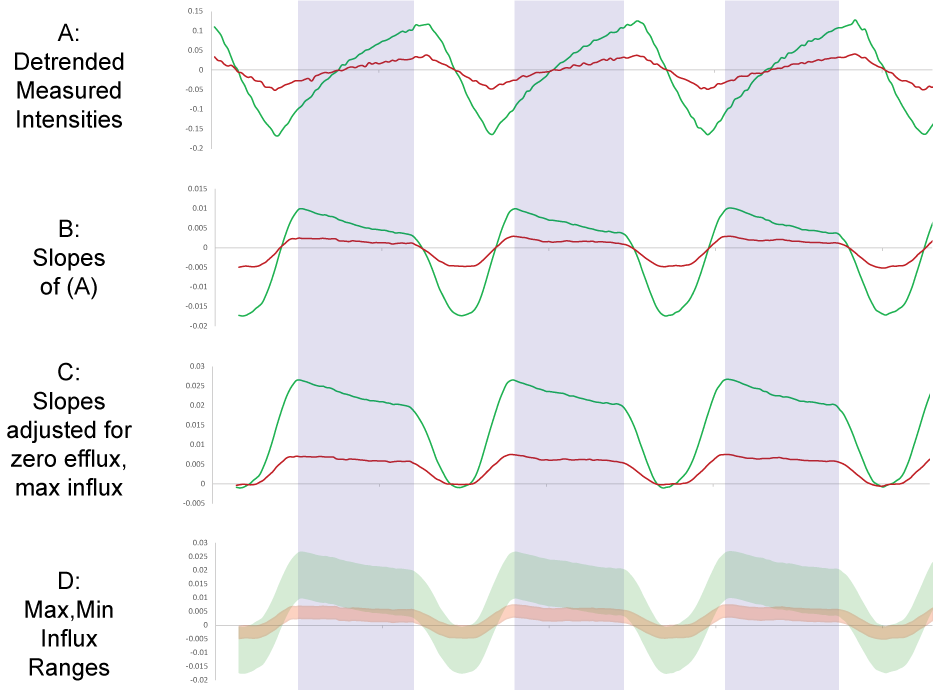
When this procedure is applied to experimental astrocyte-neuron co-culture data (**Appendix Figure 2**), the following results are obtained. While a cursory examination of the experimental traces indicates ~3-fold higher transport rates in astrocytes versus neurons, it may be argued that metabolism plays a significant role, and is greater in neurons than in astrocytes. Neuronal transport rates would then be underestimated. Using the semi-quantitative procedure described above, ranges can be derived, showing that the minimum possible influx rate of astrocytes is still larger than the maximum possible influx rate in neurons.

Appendix Figure 2: Application of theory to data

This may not, however, be the end of the story: another possibility to consider is that neurons might have a high level of resting glucose, such that the sensor is more saturated and actual neuronal glucose changes are underestimated by fluorescence changes relative to those measured in astrocytes. This cannot be the case, however, since the glucose transporters in the system are passive, such that the cytosolic glucose concentrations cannot be greater than the maximum extracellular concentrations (20 mM), at which point the response of the sensor is still approximately linear (the sensor K_d_ is ~5 mM). Even more compellingly, since the fluorescence traces oscillate stably at certain levels, the only factor that would influence tonic glucose levels would be metabolic rate. And this effect would tend to work in the opposite direction: higher metabolic rates would lead to lower tonic glucose, which in turn would tend to increase the observed magnitude of the oscillations. Accordingly, even assuming maximal glucose consumption rates in neurons (which would tend to decrease tonic glucose and increase sensor response magnitudes) and minimal consumption rates in astrocytes, the two populations are still markedly different, with astrocytes having larger glucose transport rates.

The conclusions above can be drawn directly from the current data. If, however, these results are interpreted in light of published measurements of astrocytic and neuronal metabolic rates, which show 1-2-fold greater metabolic rates in astrocytes versus neurons ^55^, the difference in the populations becomes even greater, and the factor of ~3-fold greater transport in astrocytes mentioned above is likely an underestimate.

Nevertheless, there remains a possibility that the sensor’s properties vary across different cell types, leading to different responses to equivalent glucose levels. The most likely candidate would be differences in cytosolic pH differences, which are known to affect the response of the sensor (**Supp. Fig. 2**), but published measurements reveal almost identical cytosolic pH in cultured astrocytes and neurons under similar conditions ^56^. Concentrations of other ions (K^+^, Na^+^, Ca^2+^, Mg^2+^, Cl^-^) are also similar. There may be other unknown factors perturbing sensor properties, but the above combination of data-centered reasoning and published measurements of cytological properties of cultured neurons and astrocytes support the conclusion that astrocytes in culture uptake glucose faster than neurons under identical conditions.

55. Bittner, C. X., Loaiza, A., Ruminot I., Larenas, V., Sotelo-Hitschfeld, T., Gutiérrez, R., Córdova, A., Valdebenito, R., Frommer, W. B., Barros, L. F. High resolution measurement of the glycolytic rate. *Frontiers in Neuroenergetics*. https://doi.org/10.3389/fnene.2010.00026 (2010).

56. Verkhratsky, A., Nedergaard, M. Physiology of astroglia. *Physiol. Rev.* **98**, 239-389 (2018).

**Supplementary Movie Legends**

**Supplementary Movies 1a-b. Astrocyte-neuronal co-culture appearance and responses.** At the top of each movie, the sensor channel is shown in green, and therefore includes both astrocytes and neurons. In movie 1a, neurons express the mRuby2-tagged sensor, whose red fluorescence intensity is shown over time at bottom. In 1b, astrocytes express the tagged sensor. Contrast has been adjusted to show effects more clearly; no pixels were actually saturated.

**Supplementary Movies 2a-d. Drosophila CNS explant pan-neuronal glucose responses to oscillating glucose stimulus.**

A: Green raw fluorescence channel, averaged over all planes in z.

B: Detrended data: a two-sided moving window average of size corresponding to the oscillation period was subtracted from the raw data in A, isolating the oscillating response, which travels rostro-caudally through the central neuropil and then to the peripheral cell bodies.

C: Similar to A, but with all planes of raw fluorescence data shown

D: Similar to B, but with all planes of detrended intensities shown.

**Supplementary Movies 3a-f: Zebrafish responses to various stimuli.**

A: At top, fluorescence responses of iGlucoSnFR from green channel are shown, with contemporaneous calcium (jRGECO1a) responses from the red channel shown below for comparison.

B: At top, glucose responses to blank- and insulin injections from the green channel are shown. At bottom, injections containing the tracer dye sulforhodamine 101 are shown.

C: Epinephrine responses from all planes in a volumetric confocal time lapse movie are shown as raw fluorescence. Epinephrine is added about halfway through the movie.

D: Epinephrine responses from all planes in a volumetric confocal time lapse movie are shown as ΔF/F_0_, calculated with F_0_ from immediately before epinephrine addition. Epinephrine is added about halfway through the movie.

E: Similar to D, but with ΔF/F_0_ reponses of all planes shown, with both glucose sensor from the green channel (left half) and the corresponding calcium responses from the red channel (right half).

F: All planes of brain glucose responses to a blank addition and then to an epinephrine addition to the bath are shown. Note transient reaction to both injections, but with a sustained reaction in the epinephrine addition. Also note slivers of muscle flanking posterior hindbrain which respond avidly only to epinephrine.

G: Similar to F, but with concurrent neuronal calcium responses overlaid in red. No obvious correlation was observed *prima facie.*
